# Dual targeting of salt inducible kinases and CSF1R uncouples bone formation and bone resorption

**DOI:** 10.1101/2021.02.26.433094

**Authors:** Cheng-Chia Tang, Christian D. Castro Andrade, Maureen J. Omeara, Sung-Hee Yoon, Daniel J. Brooks, Mary L. Bouxsein, Janaina da Silva Martins, Jinhua Wang, Nathanael S. Gray, Barbara M. Misof, Paul Roschger, Stéphane Blouin, Klaus Klaushofer, Annegreet Veldhuis-Vlug, Yosta Vegting, Clifford J. Rosen, Daniel J. O’Connell, Thomas B. Sundberg, Ramnik J. Xavier, Peter M.U. Ung, Avner Schlessinger, Henry M. Kronenberg, Rebecca Berdeaux, Marc Foretz, Marc N. Wein

## Abstract

Bone formation and resorption are typically coupled, such that the efficacy of anabolic osteoporosis treatments may be limited by bone destruction. The multi-kinase inhibitor YKL-05-099 potently inhibits salt inducible kinases (SIKs) and may represent a promising new class of bone anabolic agents. Here we report that YKL-05-099 increases bone formation in hypogonadal female mice without increasing bone resorption. Postnatal mice with inducible, global deletion of SIK2 and SIK3 show increased bone mass, increased bone formation, and, distinct from the effects of YKL-05-099, increased bone resorption. No cell-intrinsic role of SIKs in osteoclasts was noted. In addition to blocking SIKs, YKL-05-099 also binds and inhibits CSF1R, the receptor for the osteoclastogenic cytokine M-CSF. Modeling reveals that YKL-05-099 binds to SIK2 and CSF1R in a similar manner. Dual targeting of SIK2/3 and CSF1R induces bone formation without concomitantly increasing bone resorption and thereby may overcome limitations of most current anabolic osteoporosis therapies.

## Introduction

Osteoporosis is a major problem in our aging population, with significant health and economic burden associated with fragility fractures (1). Bone mass is determined by the balance between bone formation by osteoblasts and bone resorption by osteoclasts (2). Osteocytes, terminally-differentiated cells of the osteoblast lineage buried deep within mineralized bone matrix, sense hormonal and mechanical cues to bone and in turn regulate the activity of cells on bone surfaces (3). The majority of current osteoporosis therapeutics act by slowing down bone resorption, a strategy that most often fails to fully reverse the effects of this disease (4). Currently, bone anabolic treatment strategies are limited; development of orally-available small molecules that stimulate bone formation represents a major unmet medical need (5). Notably, efficacy of parathyroid hormone-based subcutaneous administration of osteoanabolic agents (teriparatide and abaloparatide) may be blunted by concomitant stimulation of bone resorption (6). As such, orally-available agents that stimulate bone formation without inducing bone resorption represent the ‘holy grail’ in osteoporosis drug development.

Parathyroid hormone (PTH) signaling in osteocytes stimulates new bone formation by osteoblasts (7). Salt inducible kinases (SIKs) are broadly-expressed AMPK family serine/threonine kinases (8) whose activity is regulated by cAMP signaling (9). In osteocytes, PTH signaling leads to protein kinase A-mediated phosphorylation of SIK2 and SIK3, a signaling event that suppresses cellular SIK activity (10). Genetic deletion of SIK2 and SIK3 in osteoblasts and osteocytes dramatically increases trabecular bone mass and causes phenotypic and molecular changes in bone similar to those observed with constitutive PTH receptor action (11). PTH signaling inhibits cellular SIK2/3 function; therefore, small molecule SIK inhibitors such as YKL-05-099 (12) mimic many of the actions of PTH, both *in vitro* and *in vivo* in initial studies in young, eugonadal mice (10).

Despite these advances, major unanswered questions remain regarding small molecule SIK inhibitors as potential therapeutic agents for osteoporosis. First, initial *in vivo* studies with YKL-05-099 showed increased bone formation (via a PTH-like mechanism) and, surprisingly, *reduced* bone resorption. Typically, bone formation and resorption are tightly coupled (13), and both are increased by PTH. Therefore, one goal of the current study is to define the mechanistic basis underlying the ‘uncoupling’ anti-resorptive effect of this agent. While YKL-05-099 is a potent SIK inhibitor (14), this compound also targets several other kinases (12), leaving open the possibility that some of its *in vivo* activities may be SIK-independent. Kinase inhibitor multi-target pharmacology has been exploited therapeutically for cancers whose growth is dependent on multiple activated kinases (15), yet this strategy has not been widely explored for use of kinase inhibitors in non-oncologic disease indications (16). Second, the safety and efficacy of longer-term YKL-05-099 treatment in a disease-relevant preclinical osteoporosis model remains to be determined. Finally, relevant to therapeutic efforts to develop SIK inhibitors for osteoporosis, the phenotypic consequences of post-natal SIK gene ablation are unknown.

Here we tested YKL-05-099 in female mice rendered hypogonadal by surgical oophorectomy and observed increased trabecular bone mass, increased bone formation, and reduced bone resorption. Despite these beneficial effects, toxicities of hyperglycemia and nephrotoxicity were noted. Inducible, post-natal SIK2/3 gene deletion caused dramatic bone anabolism without hyperglycemia or BUN elevation, indicating that these side effects were due to inhibition of SIK1 or other targets of YKL-05-099. Notably, inducible, global SIK2/3 gene deletion *increased* bone resorption. While YKL-05-099 potently blocked osteoclast differentiation *in vitro*, deletion of SIK2/3 or SIK1/2/3 showed no obvious effects on differentiation or function of isolated osteoclast precursors. YKL-05-099 also potently inhibited CSF1R, the receptor for the key osteoclastogenic cytokine M-CSF (17). Modeling revealed that YKL-05-099 prefers a common conformation of both CSF1R and SIK2. Consistent with these results, YKL-05-099 blocked M-CSF action in myeloid cells. Taken together, these findings demonstrate that the dual target specificity of YKL-05-099 allows this multi-kinase inhibitor to uncouple bone formation and bone resorption.

## Results

### YKL-05-099 increases trabecular bone mass in hypogonadal female mice

We previously showed that the SIK inhibitor YKL-05-099 increased bone formation and bone mass in young, eugonadal mice while simultaneously suppressing osteoclastic bone resorption (10). Based on these findings, we tested the efficacy of this compound in female mice rendered hypogonadal by surgical removal of the ovaries (OVX, Figure 1A), a common preclinical model for post-menopausal osteoporosis. In this study, 12 week-old female C57Bl/6 mice were subjected to sham or OVX surgery. 8 weeks later, mice from each surgical group were randomly divided into three treatment groups for 4 weeks total treatment. We performed side-by-side comparison of YKL-05-099 (18 mg/kg) with human PTH 1-34 (100 mcg/kg). As shown in Figure 1B-E and Supplemental Table 1, YKL-05-099 treatment increased trabecular mass bone in the femur and L5 vertebral body of hypogonadal female mice. Compared to once daily PTH (100 mcg/kg) treatment, YKL-05-099 (18 mg/kg) tended to lead to greater gains in trabecular bone mass. In contrast, this dose of PTH increased cortical bone mass and bone strength. The relationship between cortical bone mass and bone strength was preserved in response to YKL-05-099, indicating that this agent does not cause obvious defects in cortical bone quality (Figure 1F, Supplemental Table 1, Supplemental Figure 1A, B, Supplemental Table 2).

**Figure 1.**
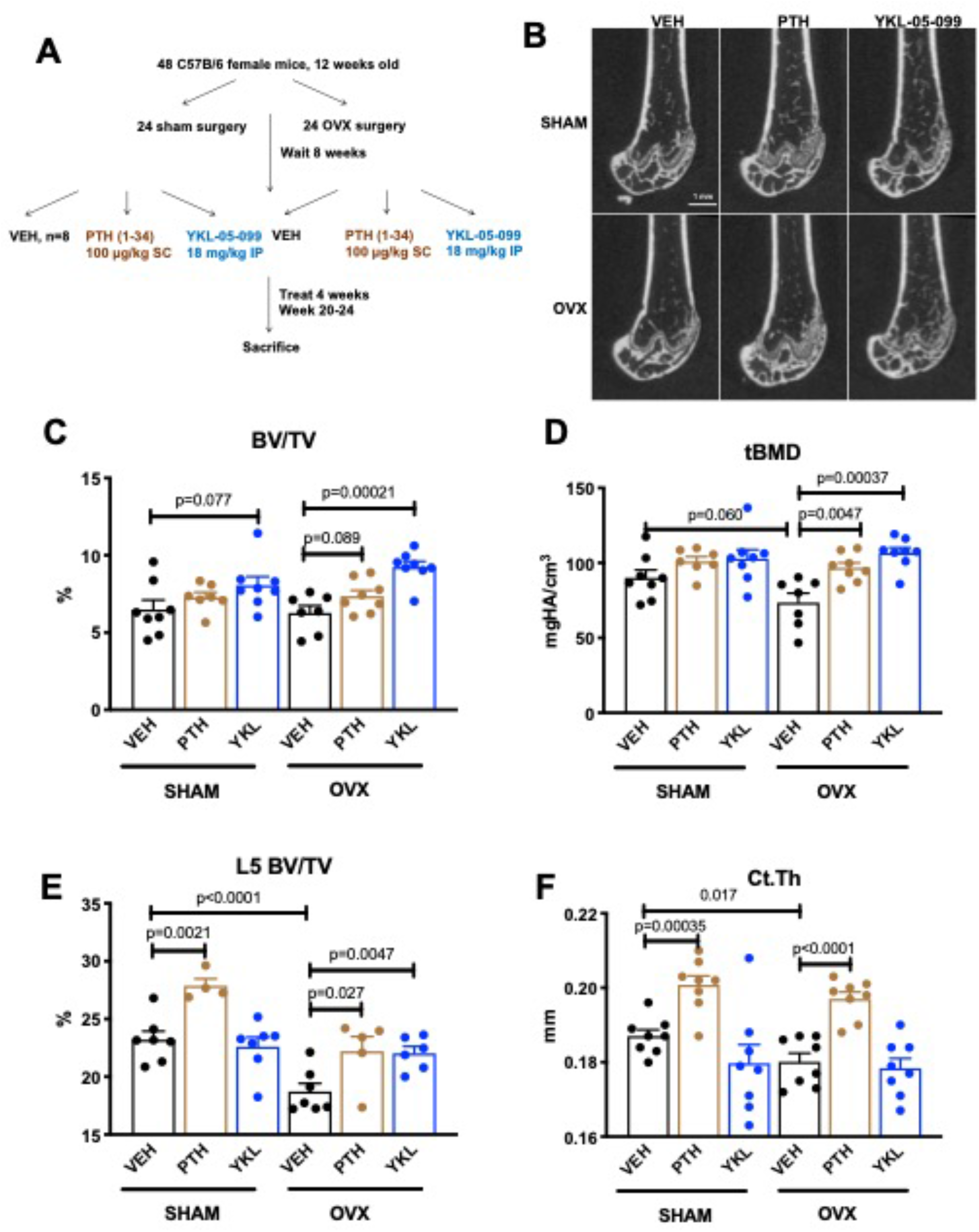
YKL-05-099 increases cancellous bone mass in hypogonadal female mice. (A) Overview of ovariectomy (OVX) study design. N=48 C57B/6 mice were subjected to sham or OVX surgery at 12 weeks of age. 8 weeks later, mice were randomly divided into the 6 indicated treatment groups, with n=8 mice per group. Animals were treated over the course of 4 weeks and then sacrificed for skeletal analyses. (B) Representative femur micro-CT images from each treatment group. Scale bar = 1 mm. (C-D) Trabecular parameters in the distal femur. BV/TV = bone volume fraction. tBMD = trabecular bone mineral density. P values between groups were calculated by one way ANOVA followed by Dunnett’s correction. All P values less than 0.1 are shown. OVX surgery reduces trabecular bone mineral density, and this is rescued by YKL-05-099 treatment. (E) Trabecular bone mass in L5. OVX surgery reduces vertebral trabecular bone mass, and this is rescued by YKL-05-099 treatment. (F) Cortical thickness in the femur midshaft. OVX surgery reduces cortical thickness. PTH (100 mcg/kg/d), but not YKL-05-099, increases cortical thickness. Also see Supplemental Table 1 for all micro-CT data from both skeletal sites. All graphs show mean ± SEM with each data point representing an individual experimental animal. A high resolution version of this figure is available here: https://www.dropbox.com/s/onfw0ued62rfsaw/Figure%201.pdf?dl=0

### YKL-05-099 uncouples bone formation and bone resorption in OVX mice

Having established that YKL-05-099 increases trabecular bone mass in OVX mice, we next sought to define the underlying cellular mechanisms. First, fasting serum was collected just prior to sacrifice to measure P1NP (a marker of bone formation) and CTX (a marker of bone resorption). Like once daily PTH treatment, YKL-05-099 treatment increased serum P1NP in both surgical groups (Figure 2A). As predicted, PTH treatment also increased bone resorption as measured by serum CTX; however, unlike PTH, YKL-05-099 treatment did not lead to statistically significant increases in serum CTX (Figure 2B). Static and dynamic histomorphometry was performed to investigate the effects of PTH and YKL-05-099 on bone cell numbers and activity at the tissue level on trabecular bone surfaces in the metaphysis of the proximal tibia. Like PTH, YKL-05-099 increased osteoblast numbers and activity (Figure 2C, E, F-H, Supplemental Table 3); however, unlike PTH (which predictably increased osteoclast numbers), YKL-05-099 tended to reduce bone resorption as assessed by histomorphometry (Figure 2D, Supplemental Table 3). Taken together, these findings demonstrate that systemic YKL-05-099 treatment boosts trabecular bone mass and bone formation as PTH does. However, unlike PTH which stimulates both bone formation and bone resorption, YKL-05-099 treatment only increases bone formation.

**Figure 2.**
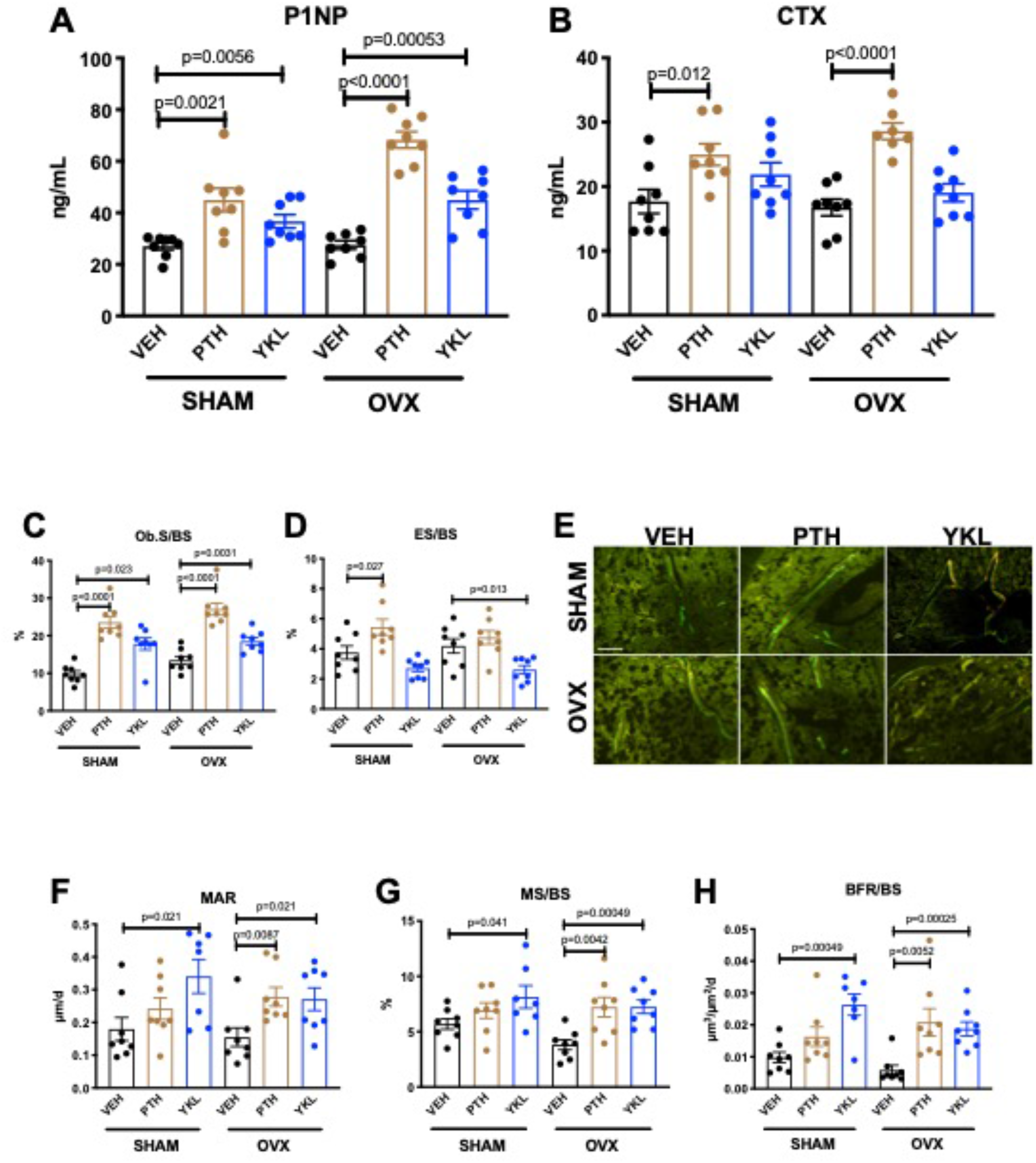
YKL-05-099 increases bone formation without increasing bone resorption in OVX mice. (A, B) Fasting serum was obtained just prior to sacrifice, following 4 weeks of treatment as indicated. P values between groups were calculated by one way ANOVA followed by Dunnett’s correction. All P values less than 0.05 are shown. Both PTH and YKL-05-099 increase levels of the bone formation marker P1NP. In contrast, only PTH treatment increases levels of the bone resorption marker CTX. P1NP levels (mean ± SD) in the different treatment groups are as follows: SHAM/VEH 27.05 (ng/ml) ± 4.07, SHAM/PTH 44.97 ± 12.9, SHAM/YKL 36.73 ± 7.33, OVX/VEH 27.53 ± 4.64, OVX/PTH 68.29 ± 9.0, OVX/YKL 44.97 ± 10.01. CTX levels (mean ± SD) in the different treatment groups are as follows: SHAM/VEH 17.69 (ng/ml) ± 5.26, SHAM/PTH 24.97 ± 4.78, SHAM/YKL 21.88 ± 5.18, OVX/VEH 16.75 ± 3.67, OVX/PTH 28.61 ± 3.4, OVX/YKL 19.04 ± 3.9. (C, D) Static histomorphometry was performed on the tibia in the proximal metaphysis to measure cancellous osteoblast surface (Ob.S/BS) and eroded surface (ES/BS). While both PTH and YKL-05-099 treatment increases osteoblast surfaces, only PTH increases bone resorption. (E) Representative fluorescent images showing dual calcein (green) and demeclocycline (red) labeling on trabecular surfaces. (F-H) Quantification of dynamic histomorphometry parameters: MAR = matrix apposition rate. MS/BS = mineralizing surface per total bone surface. BFR/BS = bone formation rate. Also see Supplemental Table 2 for all histomorphometry data. A high resolution version of this figure is available here: https://www.dropbox.com/s/gclp1oa0ubdyriw/Figure%202.pdf?dl=0

An intriguing difference between the effects of PTH and YKL-05-099 occurred at the level of osteoid surface (Supplemental Table 3). As expected, intermittent PTH treatment in OVX mice led to exuberant new bone formation leading to increased accumulation of unmineralized (osteoid) matrix on trabecular surfaces. Although YKL-05-099 increased osteoblast numbers and bone formation rate (as assessed by dual calcein/demeclocycline labeling), osteoid surface was *not* increased by this treatment. This raised the possibility that YKL-05-099 treatment might both accelerate bone matrix deposition by osteoblasts *and* its subsequent mineralization. This observation prompted us to assess bone mineralization density distribution by quantitative backscattered electron imaging (qBEI) (18) in order to assess potential effects of YKL-05-099 on bone matrix mineralization. This methodology is best suited to assess mineralization distribution patterns (BMDD) in cortical bone (where YKL-05-099 action was minimal). Only minor BMDD differences between the groups were observed (Supplemental Table 4, Supplemental Figure 1C-H). Notably, the mean calcium concentration between fluorochrome labels given 7 and 2 days prior to sacrifice (Ca_Young_, which was measured for the evaluation of Ca_Low_) did not differ greatly indicating similar mineralization kinetics between the groups based on this technique. Future study is needed to examine potential effects of YKL-05-099 on matrix mineralization in more detail.

A second, provocative effect of YKL-05-099 occurred at the level of bone marrow adipocytes (19). PTH signaling in mesenchymal lineage precursors may shift cellular differentiation from adipocyte to osteoblast lineages (20–23). As previously reported (22), acquired hypogonadism in response to OVX surgery led to increased marrow adipocytes in the metaphyseal region. YKL-05-099 treatment (Supplemental Figure 2A-C) reduced marrow adipocyte volume as assessed by semi-automated histology (24). Future studies are needed to define a potential cell intrinsic role for salt inducible kinases (or other intracellular targets of YKL-05-099) in bone marrow adipocyte differentiation and survival.

Given the clear effects observed in trabecular bone in response to YKL-05-099 treatment, we next turned our attention to the safety profile of this agent over the course of 4 weeks of treatment. Surgical treatment group (sham versus OVX) had no impact on parameters measured in vehicle-treated mice; for this reason, data are presented by drug (vehicle, PTH, or YKL-05-099) treatment. YKL-05-099 treatment had no effect on peripheral white blood cell numbers, hemoglobin, platelet counts, or absolute monocyte count (Supplemental Figure 3A). Prior to sacrifice, standard toxicology profiling was performed on fasting serum. While most parameters were unaffected by YKL-05-099 treatment, we did note mild but significant increases in blood urea nitrogen (BUN) and glucose (Supplemental Figure 3B). Hyperglycemia may be related to a potential role of salt inducible kinases downstream of hepatic glucagon signaling (25). In contrast, current genetic models do not necessarily predict nephrotoxicity from *in vivo* SIK inhibition. Taken together, these results largely demonstrate an appealing therapeutic action in bone in response to YKL-05-099 treatment. However, the potential tolerability issues observed with YKL-05-099 and key differences from the pharmacologic actions of PTH (with respect to bone resorption) prompted us to develop genetic models of adult-onset SIK isoform deletion to gain insight into whether some effects of YKL-05-099 may be related to non-SIK targets of this multi-kinase inhibitor (12).

### Inducible, global SIK2/3 deletion increases trabecular bone mass and increases bone turnover

Similar to YKL-05-099 treatment, deletion of SIK2 and SIK3 in mesenchymal lineage bone cells with Dmp1-Cre increases trabecular bone mass (11). However, unlike YKL-05-099 treatment, deletion of SIK2/3 selective in mesenchymal-lineage bone cells dramatically stimulates bone resorption. To mimic the pharmacologic effects of systemic SIK inhibitor treatment, we bred animals with ‘floxed’ SIK alleles to ubiquitin-Cre^ERt2^ mice (26) to allow global, tamoxifen-dependent SIK isoform deletion (Supplemental Figure 4A). Here, postnatal SIK3 ablation had to be postnatal to circumvent early perinatal lethality due to the key role of this kinase in PTHrP-mediated growth plate hypertrophy (11, 27). For these studies, 6 week old control (*Sik2*^f/f^ ; *Sik3*^f/f^) and SIK2/3 DKO (*Sik2*^f/f^ ; *Sik3*^f/f^ ; ubiquitin-Cre^ERt2^) mice were all treated with the same tamoxifen regimen (1 mg IP every other day, 3 injections total) to control for potential effects of tamoxifen on bone metabolism (28). Genomic DNA from cortical bone isolated two weeks after tamoxifen treatment revealed robust *Sik2* and *Sik3,* but not *Sik1*, deletion (Figure 3A). Mice analyzed three weeks after tamoxifen treatment showed overt changes in femur morphology including growth plate expansion, increased trabecular bone mass, and increased cortical porosity (Figure 3B-D, Supplemental Table 5). Growth plate histology (Figure 3E and Supplemental Figure 5) revealed expansion of proliferating chondrocytes and delayed hypertrophy, an expected phenotype in young, rapidly-growing mice.

**Figure 3.**
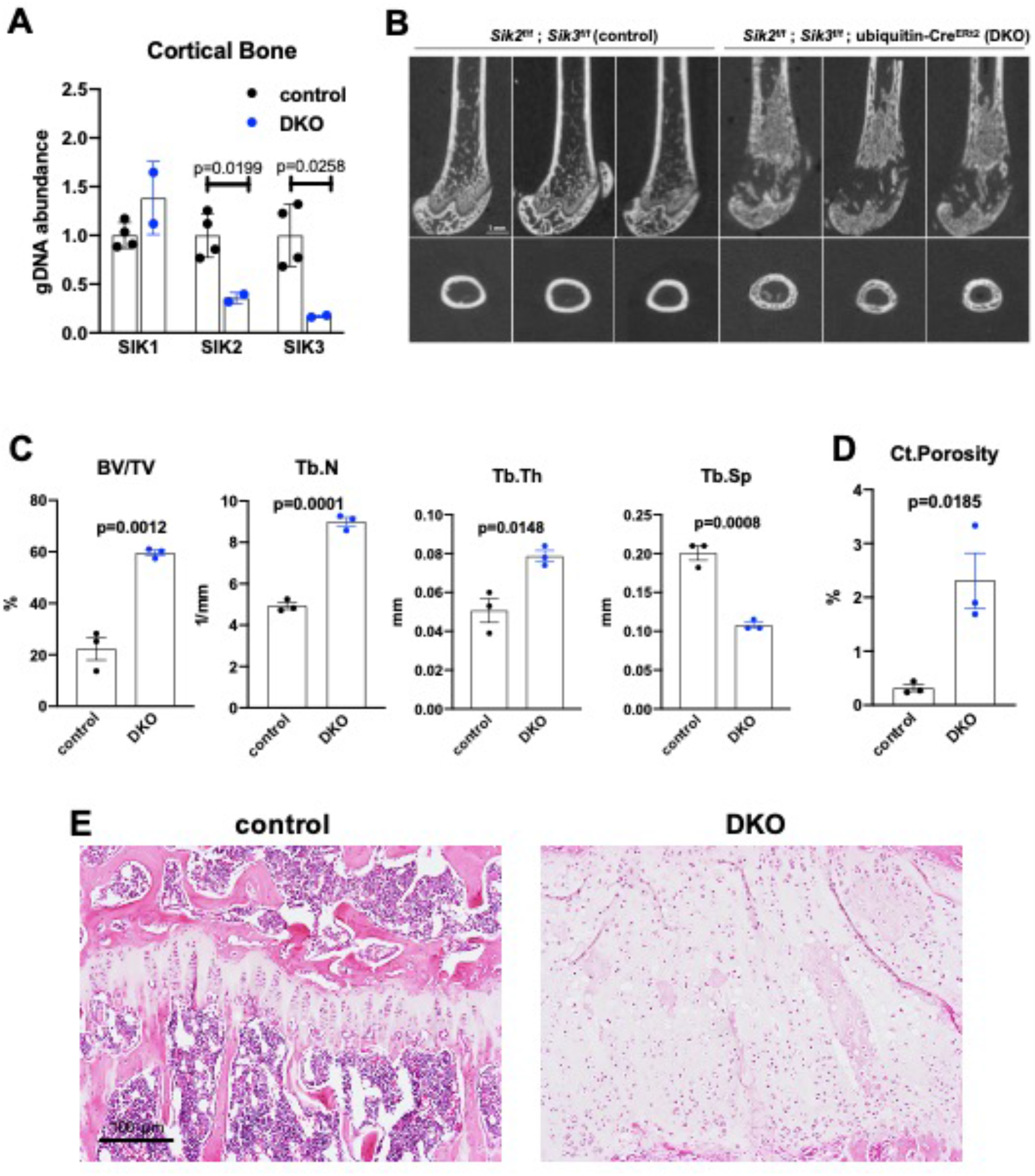
Adult onset global Sik2/3 deletion increases trabecular bone mass. (A) 6 week old *Sik2*^f/f^ ; *Sik3*^f/f^ (WT) or *Sik2*^f/f^ ; *Sik3*^f/f^ ; ubiquitin-Cre^ERt2^ (DKO) mice were treated with tamoxifen (1 mg by intraperitoneal injection, every other day, 3 doses total) and then sacrificed 14 days after the first tamoxifen injection. Cortical bone genomic DNA was isolated and SIK gene deletion was quantified. (B-D) 6 week old *Sik2*^f/f^ ; *Sik3*^f/f^ (WT) or *Sik2*^f/f^ ; *Sik3*^f/f^ ; ubiquitin-Cre^ERt2^ (DKO) mice were treated with tamoxifen (1 mg, IP, Q48H, 3 doses total) and then sacrificed 21 days after the first tamoxifen injection. Micro-CT images of the femur show increased trabecular bone mass, growth plate expansion, and increased cortical porosity. N=3 female mice of each genotype were studied. Also see Supplemental Table 5 for all micro-CT parameters measured. (E) Tibiae from mice as in (B) were stained with hematoxylin and eosin. Representative photomicrographs of the growth plate are shown. Dramatic growth plate expansion and disorganization is observed in inducible Sik2/3 mutant mice, as also demonstrated in Supplemental Figure 5. Scale bar = 100 µm. A high resolution version of this figure is available here: https://www.dropbox.com/s/mf5qxy8smrl3sg2/Figure%203.pdf?dl=0

Histology and histomorphometry from control and SIK2/3 DKO mice confirmed a dramatic increase in trabecular bone mass and an accumulation of marrow stromal cells (Figure 4A, all findings consistent with previously reported phenotypes seen when *Sik2* and *Sik3* are deleted using the *Dmp1*-Cre transgene (11)). Serum bone turnover markers (P1NP and CTX) showed increased bone formation and increased bone resorption in adult-onset, global SIK2/3 DKO animals (Figure 4B, C). Consistent with serum markers and histology, histomorphometry and TRAP (tartrate resistance acid phosphatase, osteoclast marker) staining revealed increased osteoblasts, increased osteoclasts, increased bone formation, and increased marrow stromal cell volume (Fb.V/TV) in the SIK2/3 DKOs.

**Figure 4.**
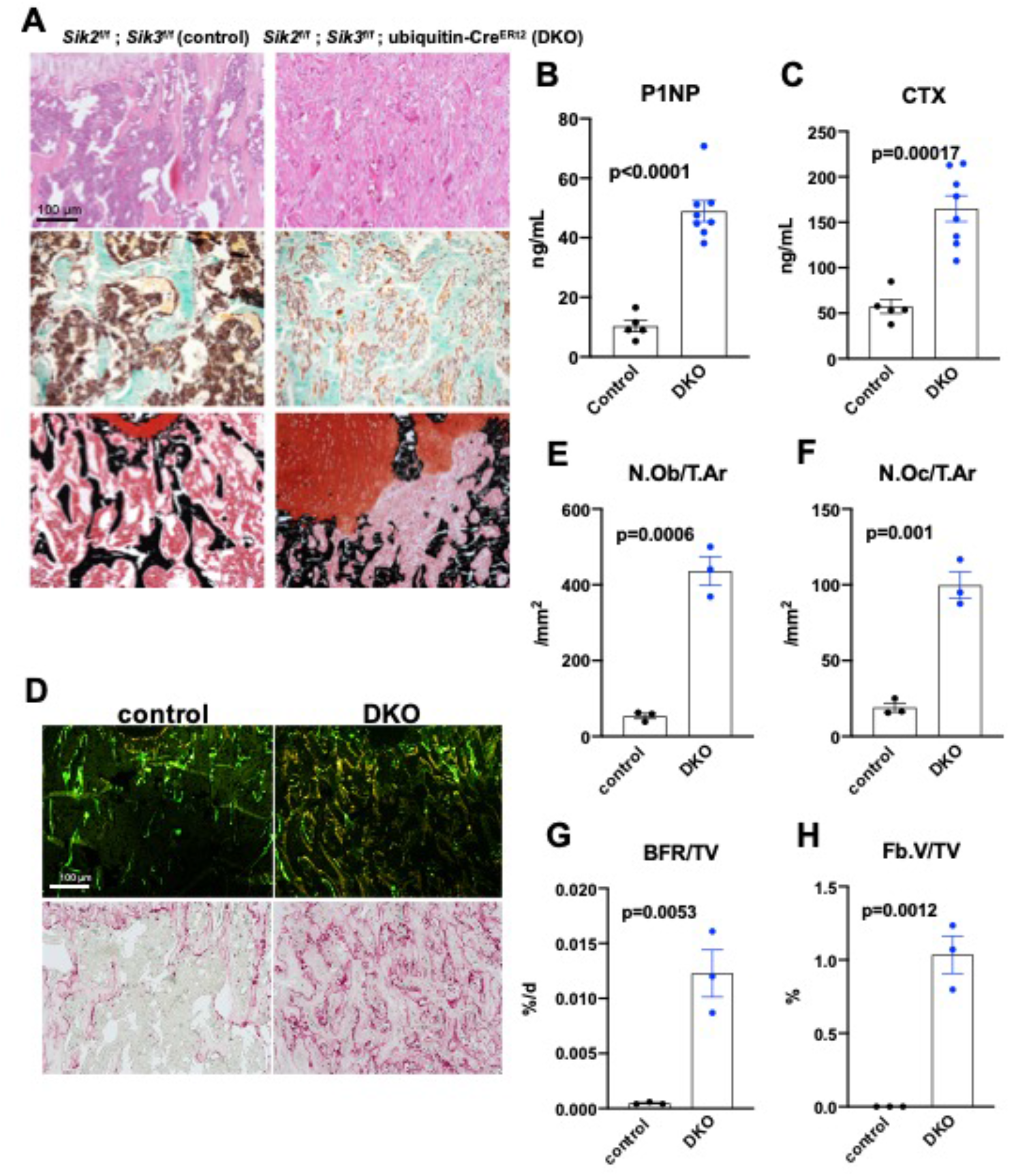
Adult onset global Sik2/3 deletion increases bone remodeling. (A) 6 week old *Sik2*^f/f^ ; *Sik3*^f/f^ (WT) or *Sik2*^f/f^ ; *Sik3*^f/f^ ; ubiquitin-Cre^ERt2^ (DKO) mice were treated with tamoxifen (1 mg by intraperitoneal injection, every other day, 3 doses total) and then sacrificed 21 days after the first tamoxifen injection. Representative photomicrographs of the proximal tibia are shown. Scale bar = 100 µm. Top panels show hematoxylin and eosin stains on decalcified paraffin-embedded sections revealing increased bone mass and marrow stromal cells in the secondary spongiosa in *Sik2/3* DKO mice. Middle panels show trichrome stains demonstrating similar findings on non-decalcified sections used for histomorphometry. Bottom panel shows dual staining with von Kossa and safranin O to demonstrate increased trabecular bone mass, expansion of marrow stromal cells, and growth plate disorganization. Also see Supplemental Figure 5 for low power images of growth plate expansion in inducible *Sik2/3* DKO mice. Results shown are representative images from n=3 female mice per genotype. (B, C) Fasting serum from mice treated as in (A) were measured for P1NP (bone formation marker) and CTX (bone resorption marker). *Sik2/3* DKO mice show increases in both bone turnover markers. (D) Top panel, mice treated as in (A) were labeled with calcein and demeclocycline at 7 and 2 days prior to sacrifice. Dark field fluorescent images show increased labeling surfaces in *Sik2/3* DKO animals. Bottom panel, decalcified paraffin sections from mice treated as in (A) were stained with TRAP (pink) to label osteoclasts. *Sik2/3* DKO mice show dramatic increases in TRAP+ cells present on bone surfaces. (E-H) Quantification of static and dynamic histomorphometry results from mice as in (A). N.Ob/T.Ar = osteoblast number per tissue area. N.Oc/T.Ar = osteoclast number per tissue area. BFR/TV = bone formation rate per tissue volume. Fb.V/TV = fibroplasia volume per tissue volume. See also Supplemental Table 6 for complete histomorphometry data. A high resolution version of this figure is available here: https://www.dropbox.com/s/6yaf398zvu0833t/Figure%204.pdf?dl=0

Prompted by the observations that YKL-05-099 treatment caused mild hyperglycemia and increased BUN (Supplemental Figure 3), similar serum toxicology profiling was performed in control and adult-onset ubiquitous SIK2/3 DKO mice. Animals were treated with tamoxifen at 6 weeks of age and then serum profiling was performed 2 weeks later. In these studies, BUN and fasting glucose levels were unaffected by global SIK2/3 ablation (Supplemental Figure 4B). Therefore, these YKL-05-099-associated toxicities are likely due to either SIK1 inhibition or off-target effects of this pharmacologic agent. These reassuring results suggest a favorable initial safety profile associated with whole body SIK2/3 gene deletion, and further support the idea of this target combination for osteoporosis drug development.

### No cell-autonomous effects of SIK gene deletion on osteoclast differentiation and function

Discordant effects at the level of bone resorption between *Sik2/3* gene deletion and YKL-05-099 treatment prompted us to study osteoclasts in more detail in this model of presumably ubiquitous inducible SIK ablation. First, we assessed ubiquitin-Cre^ERt2^ activity in myeloid osteoclast precursors by crossing ubiquitin-Cre^ERt2^ mice with tdTomato^LSL^ reporter animals. 2 weeks after *in vivo* tamoxifen treatment, >95% of bone marrow myeloid lineage cells (as marked by CD11b and LY6C expression) showed tdTomato expression (Supplemental Figure 6A), demonstrating that the ubiquitin-Cre^ERt2^ transgene is active in these cells. Bone marrow cells from control and SIK2/3 DKO mice treated with tamoxifen *in vivo* were isolated and subjected to *in vitro* osteoclast differentiation using recombinant M-CSF and RANKL (Supplemental Figure 6B). Compared to cells isolated from control mice, bone marrow cells isolated from SIK2/3 DKO mice showed increased osteoclast differentiation as assessed by increased TRAP secretion and increased numbers of TRAP-positive multinucleated cells (Supplemental Figure 6C-E). These results, consistent with evidence of increased osteoclast activity in SIK2/3 DKO mice *in vivo* yet distinct from what we observed with *in vivo* YKL-05-099 treatment, led us to further investigate a possible cell-intrinsic role of SIKs in osteoclasts.

Previous studies in murine RAW264.7 pre-osteoclastic cells suggested a potential cell intrinsic role for salt inducible kinases in osteoclast differentiation (29). Given our apparently discordant observations that (1) YKL-05-099 treatment has anti-resorptive effects and (2) SIK2/3 gene deletion increases bone resorption *in vivo,* we established a system in which SIK isoforms could be deleted *ex vivo* to test the cell-intrinsic function of SIKs in osteoclast differentiation and function. For this, bone marrow macrophages from control (*Sik2*^f/f^ ; *Sik3*^f/f^) and SIK2/3 DKO (*Sik2*^f/f^ ; *Sik3*^f/f^ ; ubiquitin-Cre^ERt2^) mice were treated with 4-hydroxytamoxifen (4-OHT) *in vitro* (30) to promote Cre^ERt2^-dependent SIK gene deletion. Using this approach with tdTomato^LSL^ ; ubiquitin-Cre^ERt2^ bone marrow macrophages, we noted that 4-OHT (0.3 µM, 72 hours) treatment led to tdTomato expression in >95% cells (Figure 5A), and robust deletion of *Sik2* and *Sik3* but not *Sik1* (Figure 5B, C). Upon M-CSF/RANKL treatment, control and SIK2/3 DKO bone marrow macrophages formed TRAP-positive multinucleated osteoclasts that were able to promote pit resorption on hydroxyapatite-coated surfaces. No significant differences were noted in osteoclast differentiation and function between control and SIK2/3 DKO cells treated with 4-OHT (Figure 5D-H).

**Figure 5.**
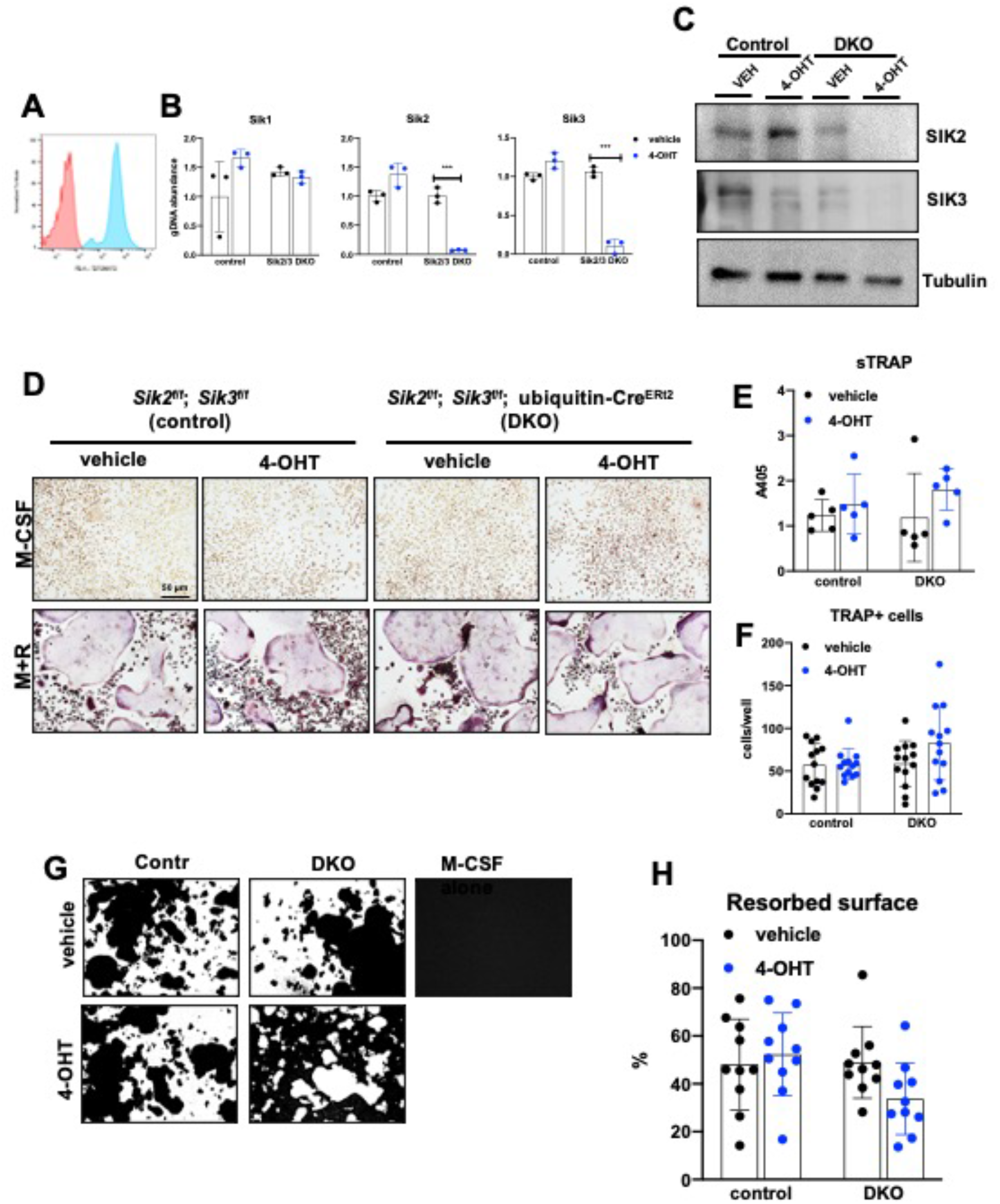
Deletion of SIK2 and SIK3 does not affect cell-autonomous osteoclast differentiation or function. (A) Bone marrow macrophages from ubiquitin-Cre^ERt2^ ; Ai14 mice were treated with vehicle (blue) or 4-hydroxytamoxifen (red, 4-OHT, 300 nM) for 72 hours followed by flow cytometry. 4-OHT treatment induces *in vitro* ubiquitin-Cre^ERt2^ activity as measured by this sensitive reporter allele. (C) Bone marrow macrophages from control or DKO mice were treated as indicated followed by immunoblotting. 4-OHT treatment leads to robust deletion of SIK2 and SIK3 protein in DKO cells. (D, E, F) BMMs from control or DKO mice were treated with vehicle or 4-OHT and then subjected to in vitro osteoclast differentiation with M-CSF plus RANKL (M+R). 4-OHT treatment in DKO cells did not cause significant changes in osteoclast differentiation as assessed by morphology (D, scale bar = 50 µm), quantifying TRAP secretion (E), or counting TRAP-positive multi-nucleated cells (F). (G, H) Osteoclasts as in (D) were grown on hydroxyapatite-coated plates in the presence of M-CSF plus RANKL. After 7 days, resorption was measured by von Kossa staining. 4-OHT treatment of DKO cells did not affect resorbed surface. M-CSF treatment alone serves as a negative control to demonstrate that pit resorption in this assay is RANKL-dependent. A high resolution version of this figure is available here: https://www.dropbox.com/s/0b9sifcjtikicd3/Figure%205.pdf?dl=0

Since YKL-05-099 inhibits all three SIK isoforms (12), the possibility remained that the inhibitory effects of this compound on osteoclast differentiation were due to SIK1 blockade. Therefore, we crossed *Sik1*^f/f^ mice (31) to *Sik2*^f/f^ ; *Sik3*^f/f^ ; ubiquitin-Cre^ERt2^ animals to create control and SIK1/2/3 triple knockouts (TKO: *Sik1*^f/f^ ; *Sik2*^f/f^ ; *Sik3*^f/f^ ; ubiquitin-Cre^ERt2^) for generation of bone marrow macrophages. Upon *ex vivo* 4-OHT treatment, TKO bone marrow macrophages showed >80% SIK gene deletion. Upon M-CSF/RANKL treatment, control and SIK1/2/3 TKO cells formed TRAP-positive multinucleated cells that promoted pit resorption. Similar to SIK2/3 DKO osteoclasts, no significant differences were noted in osteoclast differentiation and function between control and SIK1/2/3 TKO cells (Figure 6B-F). Taken together, these data argue against an important cell-intrinsic role for SIKs in osteoclasts. Moreover, these results suggest that increased bone resorption in SIK2/3 DKO mice is due to non-cell autonomous actions of SIK gene deletion, likely due to increased RANKL expression from osteoblast lineage cells similar to effects observed with PTH (10, 11, 32). Finally, these genetic data suggest that the anti-resorptive actions of YKL-05-099 may be due to engagement of intracellular target(s) other than salt inducible kinases.

**Figure 6.**
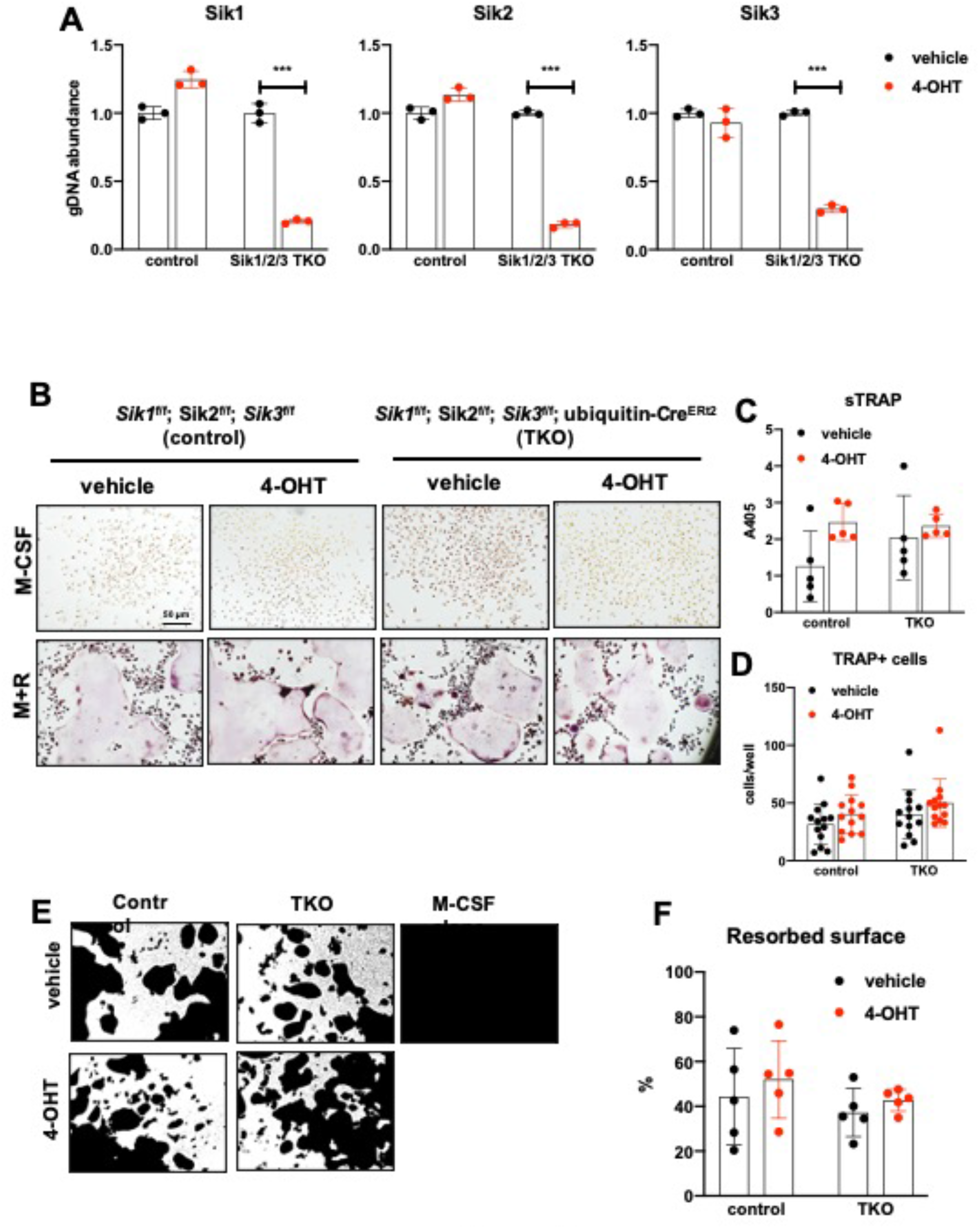
Deletion of SIK1, SIK2, and SIK3 does not affect cell-autonomous osteoclast differentiation or function. (A) Bone marrow macrophages from ubiquitin-Cre^ERt2^ ; SIK1/2/3 floxed mice were treated with vehicle (blue) or 4-hydroxytamoxifen (red, 4-OHT, 300 nM) for 72 hours. Genomic DNA was isolated for qPCR-based assessment of SIK isoform deletion. (B, C, D) BMMs from control or TKO mice were treated with vehicle or 4-OHT and then subjected to in vitro osteoclast differentiation with M-CSF plus RANKL. 4-OHT treatment in TKO cells did not cause significant changes in osteoclast differentiation as assessed by morphology (B), quantifying TRAP secretion (C, scale bar = 50 µm), or counting TRAP-positive multi-nucleated cells (D). (E, F) Osteoclasts as in (B) were grown on hydroxyapatite-coated plates in the presence of M-CSF plus RANKL. After 7 days, resorption was measured by von Kossa staining. 4-OHT treatment of DKO cells did not affect resorbed surface. M-CSF treatment alone serves as a negative control to demonstrate that pit resorption in this assay is RANKL-dependent. A high resolution version of this figure is available here: https://www.dropbox.com/s/eezv5hxawh62bl5/Figure%206.pdf?dl=0

### Modeling reveals conformation-selective preference of YKL-05-099 for SIK2 and CSF1R

Differences between YKL-05-099 treatment and *Sik2/3* gene deletion at the level of bone resorption and blood glucose and BUN prompted us to consider potential ‘off target’ kinases inhibited by YKL-05-099. First, we used TF-seq (33) to simultaneously profile the action of this compound in murine osteocytic Ocy454 cells (34, 35) at the level of 58 reporter elements that reflect output from widely-investigated signaling pathways. Using this approach, the only reporter element whose activity was significantly regulated by YKL-05-099 treatment was the CREB responsive element (Supplemental Figure 7). These findings are consistent with a known action of SIKs to regulate the activity of CRTC family CREB coactivators. When phosphorylated by SIKs, CRTC proteins are retained in the cytoplasm. Upon dephosphorylation (in this case in response to SIK inhibitor treatment), CRTC proteins translocate into the nucleus where they potentiate CREB-mediated gene expression (8, 9, 36).

TF-seq profiling of YKL-05-099 failed to inform our thinking about the anti-resorptive effects of this compound *in vivo*. However, an important clue came from review of previous data profiling YKL-05-099 binding to a panel of 468 recombinant human kinases (12). While this compound binds the active site of SIKs, it also engages a number of tyrosine kinases including the receptor tyrosine kinase CSF1R (FMS, the M-CSF receptor) (Figure 7A). Given the essential role of M-CSF action in osteoclast development (17, 37), this raised the possibility that YKL-05-099 might block osteoclast differentiation via effects on CSF1R.

**Figure 7.**
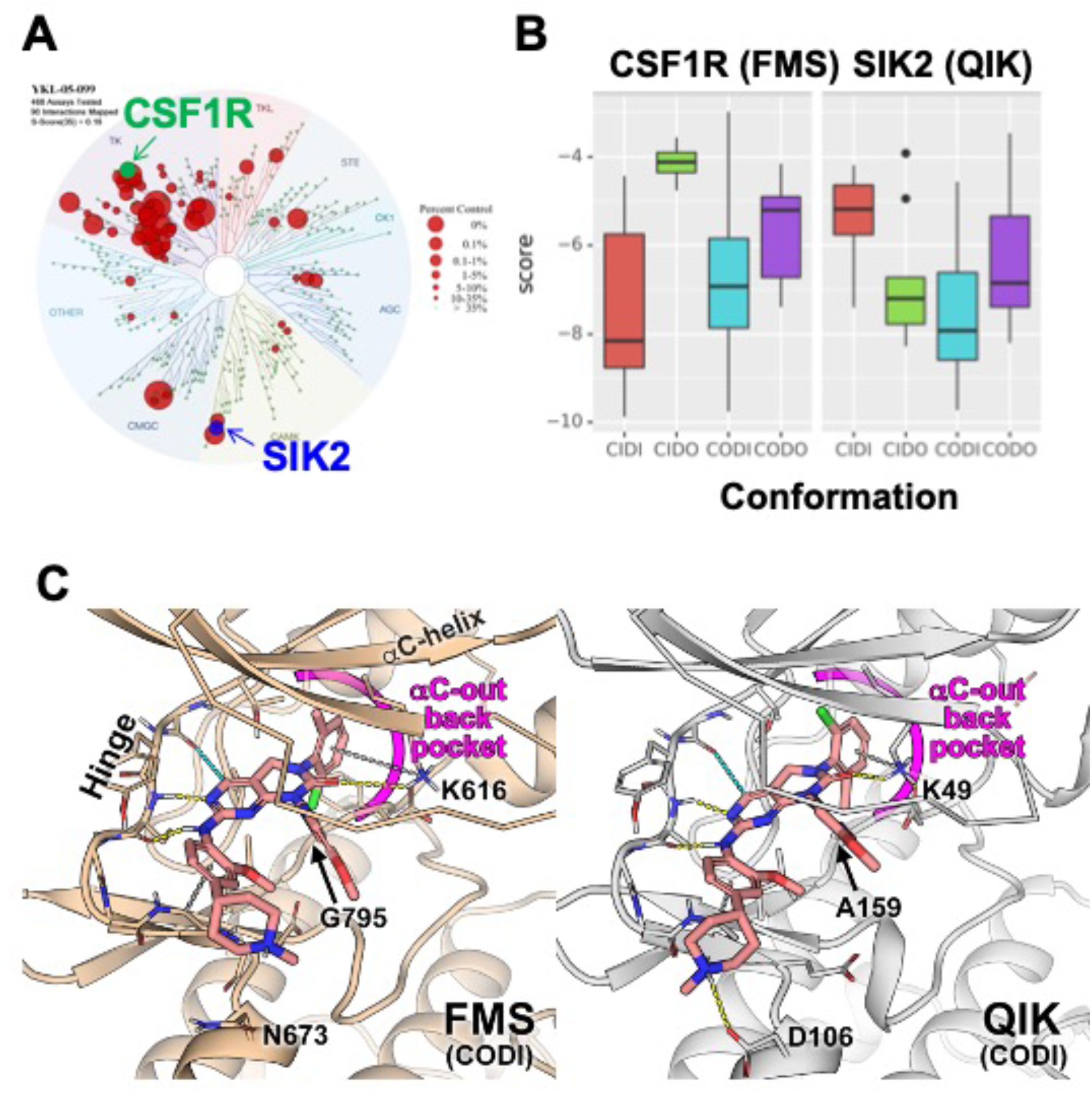
Modeling reveals YKL-05-099 preference for αC-out/DFG-in (CODI) conformation of CSF1R and SIK2. (A) Dendrogram representation of previously published (12) kinome profiling data for YKL-05-099 tested at 1.0 µM. Red circles indicate kinases with active site binding to this compound. The position of CSF1R (green) and SIK2 (blue) are noted. (B) Docking scores for YKL-05-099 binding to the active site of four kinase conformation defined by the *α*C-helix and DFG-motif for CSF1R (left) and SIK2 (right). (C) Preferential docked pose of YKL-05-099 in the modeled *α*C-out/DFG-in (CODI) conformation of CSF1R (FMS) (left) and SIK2 (QIK) (right). A high resolution version of this figure is available here: https://www.dropbox.com/s/yicizn858ee2407/Figure%207.pdf?dl=0

Most proteins within the human kinome share a highly conserved catalytic domain (38). This core domain is highly dynamic adopting a range of conformational states that are associated with catalytic activity (39–41). However, due to the limited number of atomic structures available, most kinases have only been structurally characterized in one or two conformational states (41). CSF1R (FMS) and SIK2 (QIK) represent two distinct branches of the human kinome (Figure 7A): CSF1R belongs to the receptor tyrosine kinase (TK) family, while SIKs are in the CAMK family. Although atomic structures of CSF1R have been solved in two conformational states, structure of SIKs have not been determined. Therefore, to evaluate the putative mode(s) of binding of YKL-05-099, we constructed homology models of CSF1R and SIK2 in the pharmacologically relevant conformational states.

DFGmodel (39) in combination with Kinformation (41, 42), can model four commonly observed kinase conformations defined by two critical structural elements of the kinase domain, the *α*C-helix and the DFG-motif. The four states are *α*C-in/DFG-in (CIDI), *α*C-in/DFG-out (CIDO), *α*C-out/DFG-in (CODI), and *α*C-out/DFG-out (CODO) conformations. Kinase inhibitors can also be classified by the kinase conformation they bind (43). For example, type-I inhibitors target the active CIDI conformation, whereas the type-II inhibitor sorafenib and type-I_1/2_ inhibitor erlotinib prefer the inactive CIDO and CODI conformations, respectively (41). Therefore, to model potential binding modes of a kinase inhibitor, different conformations of the kinase should be explored.

Docking of YKL-05-099 to all four modeled conformations of CSF1R and SIK2 suggested that, across both kinases, this compound shows preferences for the CODI conformation (Figure 7B). Therefore, we explored docking modes in the CODI conformation for both kinases. Three common features were noted between how YKL-05-099 engages the CODI conformation of both kinases (Figure 7C): (i) the core pyrimidine scaffold interacts with the hinge region of the kinase; (ii) the 2-chloro-6-methylphenyl moiety is buried deep in the binding pocket toward the *α*C-helix; and (iii) the 5-methoxypridin-2-yl moiety resides in the ribose binding site and the 2-methoxy-4-methylpiperdinylphenyl moiety extends outside the binding site. These modeling data demonstrate how a single kinase inhibitor can show promiscuity across unrelated kinases, and provide further support for the model that YKL-05-099 might block bone resorption via effects on CSF1R.

### YKL-05-099 blocks M-CSF action in myeloid cells

Consistent with active site binding data and modeling results, we observed that YKL-05-099 potently blocks CSF1R kinase function *in vitro* (Figure 8A, IC_50_ = 1.17 nM). Bone marrow macrophages were treated with YKL-05-099 during M-CSF/RANKL-stimulated osteoclast differentiation. In these assays, we noted potent inhibition of osteoclast differentiation in response to this compound (Figures 8B-D), with cytotoxicity noted at doses above 312 nM (Figure 8B). To more directly assess whether YKL-05-099 might block M-CSF action in pre-osteoclasts, we serum-starved cells and then re-challenged with M-CSF. M-CSF-induced receptor Y723 autophosphorylation (44) and ERK1/2 phosphorylation were completely blocked by pre-treatment with YKL-05-099 (Figure 8E). Consistent with these biochemical effects of YKL-05-099 on M-CSF receptor signaling, we also noted that YKL-05-099 pre-treatment blocked M-CSF-induced upregulation of immediate early genes *Ets2* and *Egr1* (45) (Figure 8F, G). Finally, we noted that SIK deficient osteoclasts were equally susceptible to the inhibitory effects of both the potent/selective CSF1R inhibitor PLX-5622 (46) and YKL-05-099 (Figure 8H, I, Supplemental Figures 8, 9). Taken together, these results demonstrate that YKL-05-099 can block M-CSF action in myeloid cells, serving as a likely explanation for the anti-resorptive effect seen with this agent *in vivo*.

**Figure 8.**
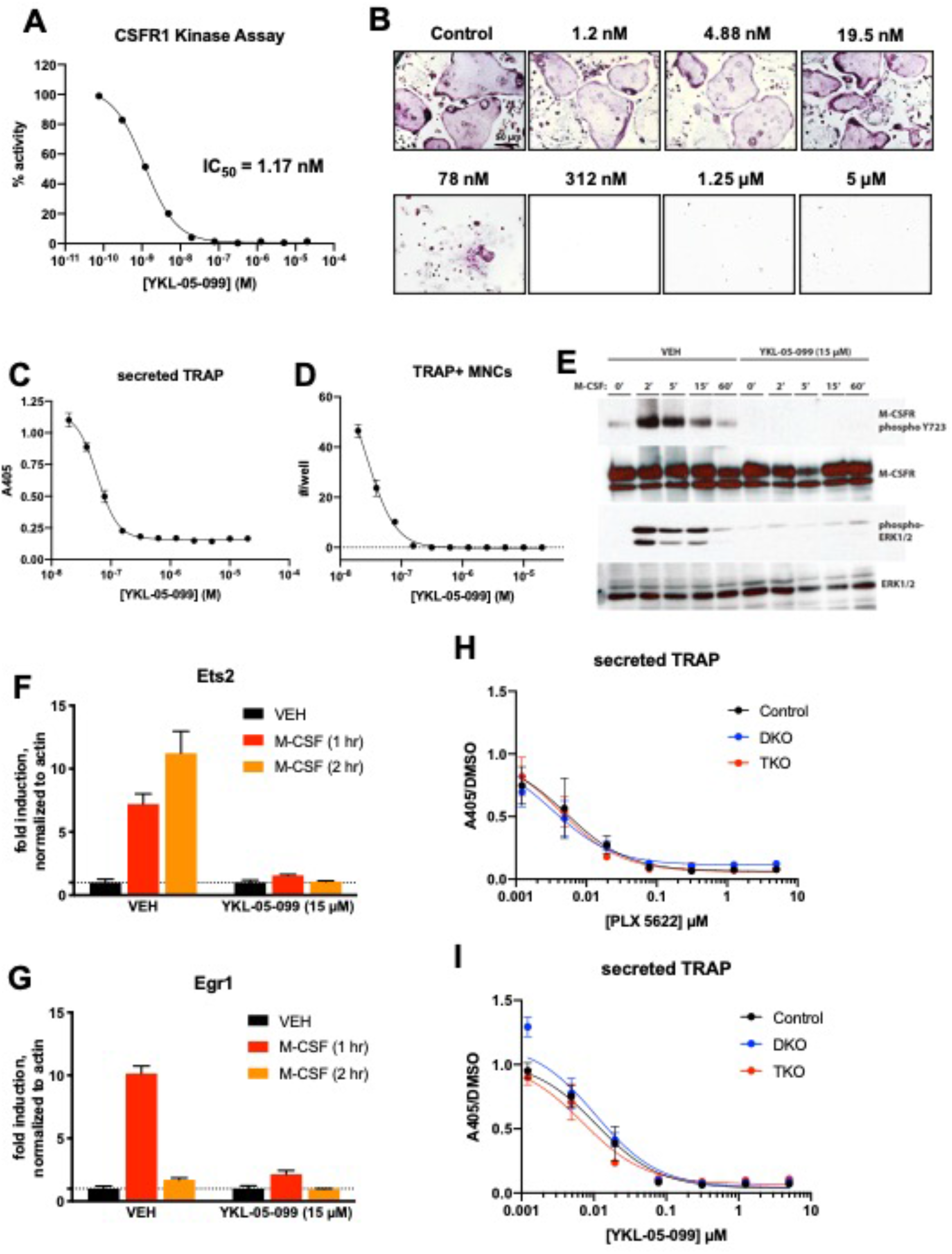
YKL-05-099 blocks M-CSF action. (A) CSF1R *in vitro* kinase assays were performed in the presence of increasing doses of YKL-05-099. This compound blocks CSF1R activity with an IC_50_ of 1.17 nM. (B) Murine bone marrow derived macrophages were grown in the presence of M-CSF, RANKL, and the indicated doses of YKL-05-099. After 3 days of differentiation, TRAP staining (purple) was performed. YKL-05-099 blocks osteoclast differentiation and causes cytotoxicity in these cultures (scale bar = 50 µm). (C) After three days of differentiation in the presence of M-CSF and RANKL, conditioned medium was collected and secreted TRAP assays were performed. YKL-05-099 treatment causes a dose-dependent reduction in TRAP secretion. Values indicate mean ± SD of n=3 wells per condition. (D) After three days of differentiation in the presence of M-CSF and RANKL, TRAP staining was performed. The number of TRAP positive multinucleated cells (MNCs) per well of a 96 well plate (n=3 wells/condition) is shown. (E) Murine bone marrow macrophages were grown in the presence of M-CSF for 5 days. Cells were then deprived of M-CSF for 6 hours, then pre-treated plus/minus YKL-05-099 (15 µM) for 60 minutes. Cells were then re-challenged with M-CSF (50 ng/ml) for the indicated times followed by immunoblotting. YKL-05-099 pre-treatment blocks M-CSF-induced M-CSFR autophosphorylation and ERK1/2 phosphorylation. (F, G) Cells as in (E) were challenged with M-CSF (50 ng/ml) for the indicated times, followed by RT-qPCR for the M-CSF target genes Ets2 and Egr1. YKL-05-099 pre-treatment blocks M-CSF-induced Ets2 and Egr1 up-regulation. (H, I) Bone marrow macrophages from ubiquitin-Cre^ERt2^ ; SIK2/3 (DKO) or ubiquitin-Cre^ERt2^ ; SIK1/2/3 (TKO) floxed mice were treated plus/minus 4-OHT as in figures 5 and 6, then plated in M-CSF/RANKL plus the indicated doses of PLX-5622 (H) or YKL-05-099 (I) for 12 days followed by secreted TRAP assays (n=6 wells from two independent experiments were assayed). In these plots, ‘control’ cells include BMMs from both DKO and TKO mice treated with vehicle prior to M-CSF/RANKL and inhibitor dose response. Irrespective of the cellular genotype, CSF1R inhibitor (PLX-5622 or YKL-05-099) treatment potently blocked osteoclast differentiation. See also Supplemental Figures 8 and 9. A high resolution version of this figure is available here: https://www.dropbox.com/s/ud8j81yh8isxad2/Figure%208.pdf?dl=0

## Discussion

New bone anabolic therapies for osteoporosis are desperately needed in order to provide more effective treatment options for this common and debilitating disease. Here we show that the small molecule kinase inhibitor YKL-05-099 boosts bone formation and trabecular bone mass in a commonly used preclinical model of post-menopausal osteoporosis. In addition to stimulating bone formation, YKL-05-099 treatment inhibits bone resorption. This appealing combination of anabolic and anti-resorptive effects is, to date, only seen with the biologic agent romosozumab (47), an anti-sclerostin antibody whose widespread use is limited due to risk of increased cardiovascular events (48).

In this study, we investigated mechanisms underlying the *in vivo* effects of YKL-05-099 treatment, and compared these results with those obtained following post-natal, ubiquitous *Sik2/3* gene deletion. While both organismal perturbations led to increased bone formation and increased trabecular bone mass, key differences were observed. First, YKL-05-099 uncoupled bone formation and bone resorption while *Sik2/3* deletion stimulated both osteoblasts and osteoclasts. As shown here, we did not observe a cell-intrinsic role for salt inducible kinases in osteoclast differentiation using *ex vivo* assays (Figures 7 and 8). Rather, YKL-05-099 likely blocks osteoclast differentiation via potent inhibition of CSF1R. As such, our current data support a model in which dual target specificity of YKL-05-099 may explain its ability to uncouple bone formation and resorption *in vivo*.

Second, YKL-05-099 caused mild hyperglycemia and increased BUN, changes not observed following *Sik2/3* deletion. Future studies are needed to better understand the mechanism of these potential (albeit mild) tolerability issues associated with YKL-05-099 treatment. Furthermore, complete characterization of metabolic and renal phenotypes in global/post-natal SIK isoform-selective and compound mutants represents a powerful future approach to better define the physiologic role of these kinases. To date, our studies have focused primarily on the function of SIKs downstream of parathyroid hormone signaling in bone (10, 11). However, potential therapeutic targeting of these kinases for the treatment of cancer, inflammation, and skin pigmentation disorders remains a high priority (9, 49). As such, the genetic tools described here to study postnatal roles of SIKs may be valuable reagents across multiple fields, in conjunction with complementary models that address the kinase function of SIK isoforms (50).

Detailed analysis of bones from OVX mice treated with YKL-05-099 for 4 weeks revealed unanticipated findings including trends towards reduced bone marrow adipocytes and potential effects on matrix mineralization. First, there may be a direct role of PTH signaling in differentiation of bone marrow adipocyte precursors (20–23). In addition, activation of other cAMP-linked GPCR signaling systems, such as ß3-adrenergic receptors, can also inhibit SIK cellular function (51, 52) and regulate bone marrow adipocyte size and numbers (53). Future studies are needed to determine whether a cell-intrinsic role exists for SIKs in bone marrow adipocyte differentiation or response to catecholamines. Second, we were surprised to note that YKL-05-099 treatment accelerated bone formation (Figure 2A, H) without increasing osteoid surface (Supplemental table 3). These findings are in stark contrast with the effects of PTH treatment which, as expected, increased bone formation and accumulation of under-mineralized bone matrix. These results suggest that YKL-05-099 somehow accelerates both matrix deposition and decreases osteoid maturation time. The assessment of Ca_Young_ (the mean calcium concentration between the double labels) which represents the calcium level at a well-defined young tissue age, did not show large differences among selected samples from all study groups. However, future studies are needed to investigate how this compound might stimulate mineralization of newly-formed bone matrix in more detail (54). Our previous studies indicated that YKL-05-099 treatment stimulates bone formation via PTH-like effects in osteocytes, including suppressing expression of the osteoblast inhibitor sclerostin (10). Whether YKL-05-099 has direct effects on osteoblast activity remains to be determined.

Achieving specificity in active site kinase inhibitors is challenging due to the conserved nature of the ATP binding pocket across protein kinases (55). It is plausible that due to small threonine residues (CSF1R: Thr663; SIK2: Thr96) at the so-called gatekeeper position (56, 57), the active sites of both CSF1R and SIK2 can accommodate multiple inhibitors (58, 59). Our modeling here further demonstrates that YKL-05-099 preferentially engages the CODI conformation of both kinases in a common binding mode. As such, multi-target binding of kinase inhibitors represents both a challenge and an opportunity as such agents are developed for chronic, non-oncologic indications (16). While avoiding undesired off-target effects will be necessary to avoid unacceptable toxicities, it remains possible that targeting precise combinations of kinases may lead to synergistic therapeutic benefits.

In summary, these findings demonstrate that a single kinase inhibitor can uncouple bone formation and bone resorption via concurrent effects on distinct kinases in distinct cell types. This work provides a framework for development of ‘next generation’ inhibitors with improved selectivity towards relevant SIK isoforms (likely SIK2 and SIK3) and CSF1R versus the remainder of the kinome. Furthermore, this work highlights the power of combining complementary genetic and pharmacologic approaches to explore the target biology and therapeutic mode of action of a small molecule kinase inhibitor.

## Materials and methods

### Mice

All animals were housed in the Center for Comparative Medicine at the Massachusetts General Hospital and all experiments were approved by the hospital’s Subcommittee on Research Animal Care. The following published genetically-modified strains were used: *Sik1* floxed mice (RRID: MGI:5648544) (31), and *Sik2* floxed mice (RRID: MGI:5905012) (25). *Sik3*^tm1a(EUCOMM)Hmgu^ mice (RRID: MGI:5085429) were purchased from EUCOMM and bred to PGK1-FLPo mice (JAX #011065) in order to generate mice bearing a loxP-flanked *Sik3* allele (11). Ubiquitin-Cre^ERt2^ mice ((26) JAX #008085) were intercrossed to *Sik1/2/3* floxed mice. In some instances, ubiquitin-Cre^ERt2^ mice were intercrossed to Ai14 tdTomato^LSL^ reporter mice (JAX #007914). Cre^ERt2^-negative littermate controls were used for all studies to account for potential influence of genetic background and impact of tamoxifen on bone homeostasis.

Both males and females were included in this study, except for the OVX studies where only female mice were used. All procedures involving animals were performed in accordance with guidelines issued by the Institutional Animal Care and Use Committees (IACUC) in the Center for Comparative Medicine at the Massachusetts General Hospital and Harvard Medical School under approved Animal Use Protocols (2019N000201). All animals were housed in the Center for Comparative Medicine at the Massachusetts General Hospital (21.9 ± 0.8 °C, 45 ± 15% humidity, and 12-h light cycle 7 am–7 pm).

For OVX studies, 12 week old sham- and OVX-operated female C57Bl/6 mice were obtained from a commercial vendor (JAX #000664). Mice from each surgical group were randomly allocated into three drug treatment groups. Drug treatments started 8 weeks after OVX surgery for a total of 4 weeks. YKL-05-099 was dissolved in PBS + 25 mM HCl and injected IP once daily five times per week for a total of 20 injections. PTH was dissolved in buffer (10 mM citric acid, 150 mM NaCl, 0.05% Tween-80, pH 5.0) and injected SC once daily five times per week for a total of 20 injections. Power calculations were performed based on previous data where eugonadal mice were treated with YKL-05-099 for two weeks (10), detailed below. For experiments in which mice were treated with either vehicle or PTH (or YKL-05-099), mice were assigned to alternating treatment groups in consecutive order. Tamoxifen (Sigma-Aldrich, St. Louis, MO, catalog #T5648) was dissolved in 100% ethanol at a concentration of 10 mg/mL. Thereafter, equal volume of sunflower oil (Sigma, catalog #88921-250ML-F) was added, the solution was vortexed and placed un-capped in a 65°C incubator overnight in order to evaporate ethanol. This working solution of tamoxifen dissolved in sunflower oil at a concentration of 10 mg/mL was used for intraperitoneal injections.

The sample size for this study was determined using the following power calculation. Grassi et al (60) performed a similar study design to assess the effects of the small molecule H_2_S donor GYY4137 on OVX-induced bone loss. In these studies, femoral BV/TV (% ± SD) fell from 7.8±2 to 4.2±1 following OVX. From these studies, a sample size of n=5/group would be required to reach 95% power to detect a “p” value of 0.05. In our published studies (10) with YKL-05-099 (6 mg/kg/d) treatment caused BV/TV to increase from 10.6±1.0 to 12.5±1.0. In these studies, a sample size of n=7/group would be required to reach 95% power to detect a “p” value of 0.05. In our preliminary data using YKL-05-099 18 mg/kg/d caused BV/TV to increase from 10.3±2.0 to 16.3±2.0. In these studies, a sample size of n=3/group would be required to reach 95% power to detect a “p” value of 0.05. In designing these experiments, a conservative estimate will be used of n=8/group, as this number reflects one additional mouse per group beyond the minimum number needed from the aforementioned scenarios. In addition, since more stringent ANOVA analysis will be required to test for an interaction between surgery and drug treatment, a larger size will be necessary. For studies investigating the effects of *Sik2/3* gene deletion on bone parameters, no statistical methods were used to predetermine sample size. The sample size was determined based on our previous experience characterizing the skeletal effects of *Sik2/3* gene deletion (11).

### Antibodies and compounds

YKL-05-099 was synthesized as previously described (12), PLX-5622 was obtained from MedChem Express (Monmouth Junction, NJ). Antibody sources and dilutions are listed below under the immunoblotting section. See Key Resources Table.

### Micro-CT

Assessment of bone morphology and microarchitecture was performed with high-resolution micro-computed tomography (μCT40; Scanco Medical, Brüttisellen, Switzerland) in 8 week old male mice. Femora and vertebrae were dissected, fixed overnight in neutral buffered formalin, then stored in 70% EtOH until the time of scanning. In brief, the distal femoral metaphysis and mid-diaphysis were scanned using 70 kVp peak X-ray tube potential, 113 mAs X-ray tube current, 200 ms integration time, and 10-μm isotropic voxel size. Cancellous (trabecular) bone was assessed in the distal metaphysis and cortical bone was assessed in the mid-diaphysis. The femoral metaphysis region began 1,700 μm proximal to the distal growth plate and extended 1,500 μm distally. Cancellous bone was separated from cortical bone with a semiautomated contouring program. For the cancellous bone region, we assessed trabecular bone volume fraction (Tb.BV/TV, %), trabecular thickness (Tb.Th, mm), trabecular separation (Tb.Sp, mm), trabecular number (Tb.N, 1/mm), connectivity density (Conn.D, 1/mm^3^), and structure model index. Transverse µCT slices were also acquired in a 500 μm long region at the femoral mid-diaphysis to assess total cross-sectional area, cortical bone area, and medullary area (Tt.Ar, Ct.Ar and Ma.Ar, respectively, all mm^2^); cortical bone area fraction (Ct.Ar/Tt.Ar, %), cortical thickness (Ct.Th, mm), porosity (Ct.Po, %) and minimum (*I*_min_, mm^4^), maximum (*I*_max_, mm^4^) and polar (*J*, mm^4^) moments of inertia. Bone was segmented from soft tissue using fixed thresholds of 300 mg HA/cm^3^ and 700 mg HA/cm^3^ for trabecular and cortical bone, respectively. Scanning and analyses adhered to the guidelines for the use of micro-CT for the assessment of bone architecture in rodents (61). Micro-CT analysis was done in a completely blinded manner with all mice assigned to coded sample numbers.

### Mechanical testing

Femora were mechanically tested in three-point bending using a materials testing machine (Electroforce 3230, Bose Corporation, Eden Prairie, MN). The bending fixture had a bottom span length of 8 mm. The test was performed in displacement control moving at a rate of 0.03 mm/sec with force and displacement data collected at 50 Hz. All bones were positioned in the same orientation during testing with the cranial surface resting on the supports and being loaded in tension. Bending rigidity (EI, N-mm2), apparent modulus of elasticity (E_app_, MPa), and ultimate moment (M_ult_, N-mm) were calculated based on the force and displacement data from the tests and the mid-diaphysis bone geometry measured with μCT. Bending rigidity was calculated using the linear portion of the force-displacement curve. The minimum moment of inertia (I_min_) was used when calculating the apparent modulus of elasticity.

### Histomorphometry

Femora were subjected to bone histomorphometric analysis. The mice were given calcein (20 mg/kg by intraperitoneal injection) and demeclocycline (40 mg/kg by intraperitoneal injection) on 7 and 2 days before necropsy, respectively. The femur was dissected and fixed in 70% ethanol for 3 days. Fixed bones were dehydrated in graded ethanol, then infiltrated and embedded in methylmethacrylate without demineralization. Undecalcified 5 μm and 10 μm thick longitudinal sections were obtained using a microtome (RM2255, Leica Biosystems., IL, USA). 5 μm sections were stained with Goldner Trichome and at least two nonconsecutive sections per sample were examined for measurement of cellular parameters. The 10 μm sections were left unstained for measurement of dynamic parameters, and only double-labels were measured, avoiding nonspecific fluorochrome labelling. A standard dynamic bone histomorphometric analysis of the tibial metaphysis was done using the Osteomeasure analyzing system (Osteometrics Inc., Decatur, GA, USA). Measurements were performed in the area of secondary spongiosa, 200 μm below the proximal growth plate. The observer was blinded to the experimental genotype at the time of measurement. The structural, dynamic and cellular parameters were calculated and expressed according to the standardized nomenclature (62).

For the adipocyte parameters, we used sections of the proximal tibia with H&E staining at 20x magnification. The following MAT outcome parameters were measured and calculated: 1] MAT volume as a percentage of the tissue volume (total adipose tissue volume: Ad.V/TV; %), 2] MAT volume as a percentage of the marrow volume (marrow adipose tissue volume: Ad.V/Ma.V; %), 3] adipocyte density (Ad.Dn; cells/mm2 marrow area) representing adipocyte number. These measurements were performed by semi-automatically tracing out individual adipocytes ‘ghosts’ in all the fields analyzed. Adipocyte ghosts appear as distinct, translucent, yellow ellipsoids in the marrow space. The total proximal tibia area below the secondary spongiosa was measured in 1-4 sections per biopsy. Adipocyte analysis was performed using a semi-automated measurement program on ImageJ (63) based image analysis software adapted from the OsteoidHisto package (64). All assessments of the sections were performed together by examiners (YV and AV-V) who were blinded to the intervention assignment.

### Serum analysis

3 hour fasting serum was collected at ZT3 from mice just prior to sacrifice by retro-orbital bleed. Serum was isolated and analyte levels were determined using the following commercially available detection kits: P1NP (IDS Immunodiagnostic Systems, #AC-33F1), CTX from IDS (#AC-06F1). All absolute concentrations were determined based on interpolation from standard curves provided by the manufacturer. DRI-CHEM 700 veterinary chemistry analyzer (Heska, Loveland, CO) was used for measurement of serum analytes: albumin, alkaline phosphatase, ALT, BUN, calcium, cholesterol, globulin, glucose, phosphorus, total bilirubin, triglycerides, and total protein. For complete blood counts, whole blood was collected into heparin-coated tubes and kept on ice. CBCs were measured on a 2015 Heska Element HT5 Veterinary Hematology Analyzer. HeskaView Integrated software was used for data analysis.

### Quantitative Backscattered Electron Imaging (qBEI)

Distal femora from 6 study groups were analyzed (n=8 each group): SHAM VEH, SHAM PTH, SHAM YKL, OVX VEH, OVX PTH, and OVX YKL. These bones were embedded undecalcified in polymethylmethacrylate (PMMA) and measured for bone mineralization density distribution (BMDD) using qBEI. The surfaces of the sample blocks were flattened by grinding and polishing (Logitech PM5, Glasgow, Scotland) and carbon coated so as to facilitate qBEI. A scanning electron microscope equipped with a four quadrant semiconductor backscatter electron detector (Zeiss Supra 40, Oberkochen, Germany) was used. Areas of metaphyseal (MS) and cortical midshaft bone (Ct) were imaged with a spatial resolution of 0.88 µm/pixel. The gray levels, reflecting the calcium content, were calibrated by the material contrast of pure Carbon and Aluminum. Thus, the resulting gray level histograms could be transformed into calcium weight percent (wt% Ca) histograms (Supplemental Figure 1D) as described previously (65). Five parameters were derived to characterize the BMDD (18). For information about Ca_Young_, which is the mean calcium concentration of the bone area between the double fluorescence labels, the identical bone surface measured with qBEI was additionally imaged in a Confocal Laser Scanning Microscope (Leica TCS SP5, Leica Microsystems CMS GmbH, Wetzlar, Germany) using a laser light of 405 nm for fluorescence excitation and a 20x object lens (pixel resolution of 0.76 µm). By matching the CLSM with the qBEI images, the sites of the fluorescence labels were overlaid exactly onto the qBEI images (Supplemental Figure 1E). Ca_Young_ was obtained from a total of 39 areas from a subgroup of nine samples and was subsequently used to calculate Ca_Low_, which reflects the percentage of newly formed bone area (Suppl. Fig. 1H).

### Flow cytometry

Ubiquitin-Cre^ERt2^ ; tdTomato^LSL^ mice were treated with tamoxifen (1 mg IP Q48H, three injections total). Two weeks after the first tamoxifen injection, mice were sacrificed and bone marrow cells were isolated by flushing with ice cold PBS using a 25G needle. 10^6^ bone marrow cells were protected from light and stained on ice for 30 minutes with the following primary antibodies in a 10 µL staining volume. APC anti-mouse/human CD 11b antibody (Biolegend, San Diego, CA), FITC anti-mouse CD3 antibody (Biolegend, San Diego, CA), and PE/Cy7 anti-mouse Ly-6C antibody (Biolegend, San Diego, CA). After staining, cells were washed twice with FACS buffer (PBS plus 2% heat inactivated fetal bovine serum) and analyzed on a SORP 8 Laser BD LSR flow cytometer (Becton, Dickinson and Company, Franklin Lakes, NJ).

### Bone marrow macrophages and osteoclast differentiation

Bone marrow cells were harvested from 4 week-old C57BL/6 mice. Femur and tibia were removed and the marrow cavity was flushed out with 10mL 1x phosphate-buffered saline per mouse (GE Healthcare Life Sciences, MA) with a 25 G needle (Becton, Dickinson and Company, Franklin Lakes, NJ)) which was then centrifuged at 1,000rpm for 10 minutes. Supernatant was removed, and cell pellet was resuspended in 1mL red blood cell lysis buffer (Sigma-Aldrich, St. Louis, MO) and incubated for 2 minutes followed by centrifugation at 1,000rpm for 5 minutes to collect cells. Bone marrow cells were plated on 100mm non-treated tissue culture dishes (Corning Inc, Corning, NY) and maintained in alpha minimum essential medium eagle (MEM) (Sigma-Aldrich, St. Louis, MO) containing 15% fetal bovine serum FBS (Gemini Bio, West Sacramento, CA), 1x penicillin-streptomycin (Gibco, Waltham, MA), 1x GlutaMAX supplement (Gibco, Waltham, MA) and in the presence of 30ng/mL recombinant mouse M-CSF (R&D systems, Minneapolis, MN). These cells were maintained at 37°C in a humidified 5% CO_2_ incubator for 3 days, then collected by trypsinization. Primary osteoclast precursors were then seeded in 96 well plates at 2,000 cells/well (20,000 cells/mL), and differentiation was induced with 50ng/mL of recombinant mouse RANKL (R&D systems, Minneapolis, MN) and 30ng/mL recombinant mouse M-CSF (R&D systems, Minneapolis, MN). Fresh medium was replenished every 3 days.

For experiments with 4-hydroxytamoxifen treatment, primary osteoclast precursors were seeded after initial culture on non-treated plates into 6 well plates in the presence of 30ng/mL recombinant mouse M-CSF (R&D systems, Minneapolis, MN) for 3 days. Cells were then treated with 300nM of 4-hydroxytamoxifen (Sigma-Aldrich, St. Louis, MO) or vehicle (methanol, Sigma-Aldrich, St. Louis, MO) for 3 days. Fresh medium with 30ng/mL recombinant mouse M-CSF (R&D systems, MN) was then added for another 3 days prior to collecting cells for subsequent osteoclast differentiation.

For osteoclast differentiation assays, primary osteoclast precursors were seeded in 96 well plates, and differentiation was induced with 50ng/mL of recombinant mouse RANKL (R&D systems, Minneapolis, MN) and 30ng/mL recombinant mouse M-CSF (R&D systems, Minneapolis, MN). Fresh medium was replenished every 3 days. sTRAP assay was measured with acetate buffer from acid phosphatase leukocyte kit (Sigma-Aldrich, St. Louis, MO), 1M sodium L-tartrate dibasic dihydrate (Sigma-Aldrich, St. Louis, MO) and pNPP substrate (Sigma-Aldrich, St. Louis, MO). 50µL of cell culture supernatant was transferred to a new 96 well plate and 150µL of substrate mix was added. Plate was then incubated 37°C for an hour. Reaction was terminated by adding 3N NaOH (Sigma-Aldrich, St. Louis, MO), which results in an intense yellow color. Absorbance was measured at 405nm.

TRAP staining was performed using acid phosphatase leukocyte kit (Sigma-Aldrich, St. Louis, MO) to visualize osteoclasts. TRAP buffer was prepared freshly on the day of experiment by mixing acetate buffer containing sodium acetate and acetic acid with sodium tartrate and naphthol AS-BI phosphate disodium salt (Sigma-Aldrich, St. Louis, MO). TRAP staining solution was prepared by mixing Fast Garnet GBC Base with sodium nitrite. For TRAP staining, culture medium was removed from cells, and cells were fixed with fixative solution for 5 minutes at room temperature. Fixative solution was prepared fresh on the day of experiment by mixing citrate solution, acetone with 37% formaldehyde. Cells were then washed twice with deionized water and stained with TRAP staining solution for 60 minutes at 37°C.

For pit resorption assays, primary osteoclast precursors were seeded in 96 well resorption plates at 2000 cells/well (Corning, Inc, Corning, NY), and differentiation was induced with 50ng/mL of recombinant mouse RANKL and 30ng/mL recombinant mouse M-CSF. Fresh medium with M-CSF and RANKL was replenished every 3 days. Culture was aspirated and cells were washed three times with 1x PBS. 100µL of 10% bleach was added to each well and incubated for 30 minutes at room temperature. Bleach solution was removed and cells were washed twice with deionized water. Plates were air-dried at 4C overnight and osteoclast resorption pits were visualized the next day. Pit resorption was quantified using ImageJ.

### gDNA deletion analysis

Genomic DNA was isolated from cultured bone marrow macrophages or cortical bone using DNeasy Blood & Tissue Kit (Qiagen, Hilden, Germany) according to the instructions of the manufacturer. For cultured bone marrow macrophages, cell were treated with vehicle (methanol) or 4-hydroxytamoxifen (0.3 µM) for 72 hours prior to gDNA isolation. For cortical bone, mice were treated with tamoxifen (three 1 mg IP injection every 48 hours) and then sacrificed 7 days after the first tamoxifen injection. Epiphyses were removed and bone marrow cells were flushed using ice cold PBS. Remaining cortical bone fragments were flash frozen in liquid nitrogen and then pulverized using a morter and pestle. Bone tissue was then subjected to gDNA isolation and quantified using a spectrophotometer (Nanodrop 2000, Thermo Fisher Scientific, Waltham, MA). 15 ng gDNA was used for each qPCR reaction. For each gene, one primer pair was used that was internal to the targeted loxP sites and a second external primer pair was used to normalize to input gDNA amount. Relative abundance of the targeted gene was calculated by the 2(-Delta Delta C(T)) method using the external primer pair as control (66).

### TF-seq

Ocy454 cells were infected with lentiviral particles expressing TF-seq library (33) and eGFP. Infected cells were isolated by sorting for GFP expression. The cell line was confirmed to be mycoplasma-free by PCR. Thereafter, cells were plated at a density of 50,000/ml in 96 well plates (5,000 cells/well) and allowed to expand at 33°C for 48 hours. Cells were then moved to 37°C for 24 hours, then treated with vehicle (DMSO) or YKL-05-099 for time points ranging from zero minutes to 4 hours with n=3 wells per condition. Cells were washed with ice cold PBS and then lysed with RLT buffer (Qiagen, supplemented with ß-mercaptoethanol) and stored at −80°C. RNA was purified using Agentcourt RNA Clean XP to precipitate the nucleic acids in 1.25 M NaCl and 10% PEG-8000. Maxima reverse transcriptase (Thermo Fisher Scientific) was run according to the manufacturer’s instructions using a multiplexed primed reverse transcriptase reaction. The biotinylated 96-well sequence tagged TF-seq-specific reverse transcriptase primers, and the biotinylated 96-well degenerate sequence-tagged polydT reverse transcriptase primers were used at 750 and 250 nM final concentrations, respectively, with 50 units of Maxima. After sequencing-tagging all cDNA during reverse transcriptase, each 96-well plate was pooled and the unincorporated primers were washed away from the cDNA by precipitating with 10% PEG-8000 and 1.25 M NaCl. Amplification of the TF-seq gene reporter amplicon was performed on 50% of the cDNA using primers with full Illumina-compatible sequencing adapters in a 600µl PCR reaction for 28 cycles. The 422-bp amplicon was then gel extracted for sequencing. TF-seq is a 50-bp single-end read, well-tag, RNA UMI, followed by a 17-bp constant sequence, then the reporter tag UMI, and ultimately the reporter tag. We counted the number of unique RNA molecules for every well and reporter element, requiring a perfect match for the respective tags. UMI tag counts for each reporter element were obtained. Reporter element activity was expressed as fold change versus baseline (time 0) for each drug treatment.

### Generation of models of kinases in various conformational states

Models of CSF1R (FMS), SIK1, and SIK2 (QIK) in four conformations, the CIDI, CIDO, CODI, and CODO states, were generated using DFGmodel (39). DFGmodel uses a multi-template homology modeling approach to construct composite homology models in various kinase conformations. The models are based on an augmented version of a structurally validated sequence alignment of human kinome (67). For the CIDO, CODI, and CODO states, DFGmodel uses multi-template approach, where manually curated sets of determined structures, covering a unique range of conformations for each state are used as template structures. For the CIDI, DFGmodel uses a single-template approach to construct the models; DFGmodel searches a library of kinase structures annotated by Kinformation (41, 42, 68) (www.kinametrix.com) to identify a best-matched kinase in CIDI state as template. DFGmodel then uses MODELLER (69) v9.21 to generate homology models for each kinase. For CIDI, CIDO, and CODO states, 50 models were generated; POVME (70) v2.1 was used to estimate the binding site volume of each model, where 10 models with the largest volume were selected. For CODI, 4 sets of templates are required to cover the observed conformation, thus 40 models were generated for each of these sets; 8 models with the largest volume in each of the sets were selected. Lastly, CSF1R crystal structures are also included for examination.

### Molecular docking

Molecular docking was performed using Glide (71) (Schrödinger 2019-3). Default settings and Standard Precision (SP) model, with the addition of aromatic hydrogen and halogen bonds, were used for Glide protein grid generation and docking of inhibitor into the active site of protein kinase models in CIDI, CIDO, CODI, and CODO conformations. OPLS3e force field (72) was used to parameterize both protein and ligands. Scaling of van der Waals (vdW) radius for receptor atoms was set to 0.75. The docking results were averaged for 10 models, as described by Ung et al. (39). The small molecule YKL-05-099 was prepared with Schrödinger’s LigPrep program. Docking pose of this molecule in each of the models was examined; poses that do not possess the critical hydrogen-bonds with the hinge residues or have docking score higher than −7.0 were rejected.

### CSF1R kinase assay

Assays were performed in base reaction buffer (20 mM Hepes (pH 7.5), 10 mM MgCl2, 1 mM EGTA, 0.02% Brij35, 0.02 mg/ml BSA, 0.1 mM Na3VO4, 2 mM DTT, 1% DMSO). YKL-05-099 was dissolved in 100% DMSO in a 10 mM stock. Serial dilution was conducted by Integra Viaflo Assist in DMSO. Recombinant CSF1R (Invitrogen, Carlsbad, CA) was used at a concentration of 2.5 nM. The substrate used was pEY (Sigma-Aldrich, St. Louis, MO) at a concentration of 0.2 mg/ml. Kinase assays were supplemented with 2 mM Mn^2+^, and 1 µM ATP was added. Assays were performed for 20 minutes at room temperature, after which time ^33^P-ATP (10 µCi/µl) was added followed by incubation for another 120 minutes at room temperature. Thereafter, radioactivity incorporated into the pEY peptide substrate was detected by filter-binding method. Kinase activity data were expressed as the percent remaining kinase activity in test samples compared to DMSO reactions. IC_50_ values and curve fits were obtained using Prism 8.0 GraphPad Software (GraphPad Software, San Diego, CA).

### Immunoblotting

Immunoblotting was performed in lysates derived from primary bone marrow macrophages. Cells were scraped into ice cold PBS and cell pellets were lysed in TNT (200 mM NaCl, 20 mM Tris HCl pH 8, 0.5% Triton X-100 supplemented with protease and phosphatase inhibitor cocktail (Thermo Fisher Scientific, Waltham, MA). Cells were vortexed twice in lysis buffer for 30 seconds followed by centrifugation at top speed at 4C for 5 minutes. Protein concentration in cell lysates was quantified using Bradford assay (Thermo Fisher Scientific, Waltham, MA). Equal amounts of protein were then separated by SDS-PAGE under reducing conditions. Proteins were then transferred to nitrocellulose membranes. Membranes were blocked in TBST plus 5% milk for 30 minutes at room temperature. Thereafter, membranes were incubated overnight at 4C in TBST with 5% BSA plus primary antibodies (source and dilutions below). Membranes were then washed three times with TBST for 5 minutes, followed by secondary antibody incubation (1:2000 goat anti-rabbit) for 30 minutes at room temperature. Next, membranes were washed again for three times with TBST for five minutes followed by detection with Pierce^TM^ ECL Plus Western Blotting Substrate (Thermo Fisher Scientific, Waltham, MA) and imaging by Azure c600. The primary antibodies were: SIK2 (1:1000), CSF1R phospho-Y723 (1:1000), CSF1R (1:1000), phospho-ERK1/2 (1:1000), ERK1/2 (1:2000), Tubulin (1:1000) were obtained from Cell Signaling Technology (Danvers, MA). Anti-SIK3 rabbit polyclonal antibodies (1:1000) were obtained from Abcam (Cambridge, United Kingdom).

### Statistical analysis

When two groups were compared, statistical analyses were performed using unpaired two-tailed Student’s t-test. When more than two experimental groups were compared, one-way ANOVA (GraphPad Prism 8.0) with Dunnett’s correction was performed. P values less than 0.05 were considered to be significant. The numbers of mice studied in all experiments are described in figure legends, and in all figures data points represent individual mice. All data points indicate individual biologic replicates (independent experimental samples) and not technical replicates (the same sample re-analyzed using the same method). A Source Data File is presented for all figures.

## Acknowledgements

We thank Drs. Michael Mannstadt, Lauren Surface, Francesca Gori, Tatsuya Kobayashi, Mark Poznansky, Christiana Iyasere, and members of the Wein laboratory for helpful discussions. MNW acknowledges funding support from the American Society of Bone and Mineral Research, the Harrington Discovery Institute, the MGH Department of Medicine Innovation Program, and the National Institute of Health (DK116716, AR066261, and AR067285). HMK acknowledges funding support from the National Institute of Health (AR066261 and DK011794). MF acknowledges funding support from Centre National de la Recherche Scientifique (CNRS), the Société Francophone du Diabète (SFD) and the Fondation pour la Recherche Médicale (FRM). RB acknowledges funding support from the National Institutes of Health (DK092590 and AR059847).

---

**Supplemental Figure 1.**
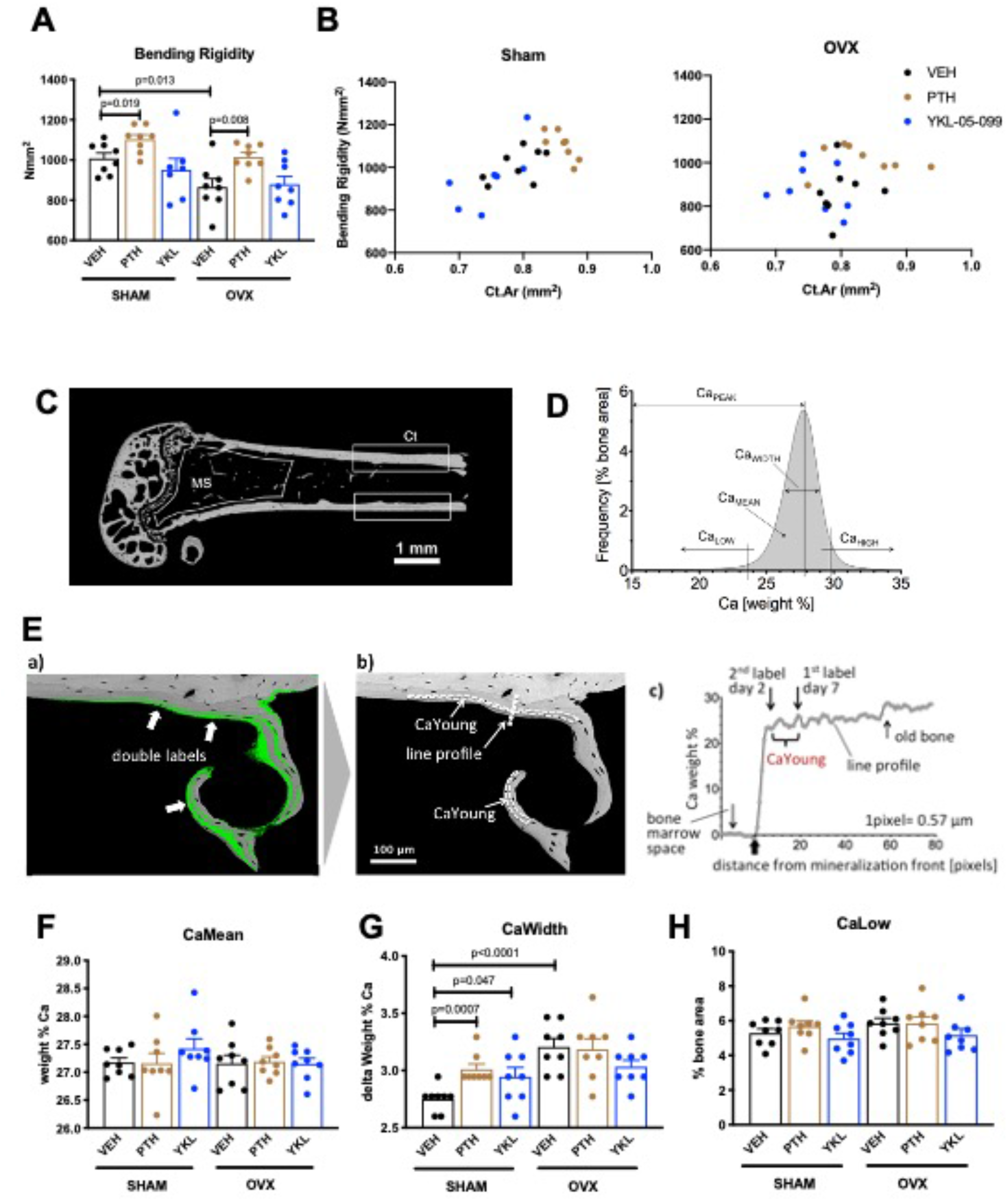
Results of biomechanical testing (A,B) and quantitative backscattered electron imaging (qBEI, C-H) from OVX study. (A) Bending rigidity results. Consistent with cortical bone micro-CT data, only PTH treatment increases femur mechanical strength. P values between groups were calculated by one way ANOVA followed by Dunnett’s correction. All p values less than 0.05 are shown. See Supplemental Table 3 for additional parameters obtained from three point bending testing. (B) The relationship between cortical bone mass (x-axis, Cortical Area) and bone strength (y-axis, Bending rigidity) was plotted for individual mice in the indicated surgical and drug treatment groups. No obvious changes in the bone mass/strength relationship were noted in response to PTH or YKL-05-099 compared to vehicle-treated animals. (C) qBEI was performed on distal femur from OVX study mice (n=8 per each group) to determine the effects of OVX and drug treatments on bone mineralization in metaphyseal spongiosa (MS) and midshaft cortical bone (Ct). (D) BMDD from a representative SHAM-VEH mouse in the cortical midshaft region. The BMDD parameters are depicted: Ca_Mean_ (mean Ca content), Ca_Peak_ (mode Ca content), Ca_Width_ (the full width at half maximum of the distribution), Ca_Low_ (the percentage of mineralized bone with a calcium concentration less than 23.84 weight % corresponding to Ca_Young_), and Ca_High_ (the percentage of bone areas with a calcium concentration beyond the 95^th^ percentile of the SHAM-VEH group BMDD, which was 29.81 weight % Ca in cortex and 28.25 weight % Ca in the metaphysis). (E) For the evaluation of Ca_Low_, the mean calcium concentration between the two fluorescence labels Ca_Young_ (corresponding to a mineralized tissue age of 2 to 7 days) was determined. For this purpose, the (a) qBEI image was overlaid with a matched confocal scanning laser microscope image (fluorescence mode) from the identical sample surface, and (b) Ca_Young_ was measured in the bone areas between the double labels (areas enclosed by dotted lines). (c) Example of mineralization of a line profile through an area of new bone formation as indicated in (b), showing the rapid primary increase and second slower increase in bone matrix mineralization. Ca_Young_ represents the calcium level at the beginning of the second phase. Graphs in (F-H) show the effects of surgery and drug treatment on cortical bone Ca_Mean_, Ca_Width_ and Ca_Low_ (for cortical bone Ca_Peak_ and Ca_High_ as well as metaphyseal BMDD results, see Supplemental Table 4). In sham-operated mice, PTH treatment increases Ca_Width_. In general, minimal effects of surgery or drug (PTH or YKL-05-099) were observed as assessed by qBEI. A high resolution version of this figure is available here: https://www.dropbox.com/s/bq0g6jc15eyhnfl/Figure%20S1.pdf?dl=0

**Supplemental Figure 2.**
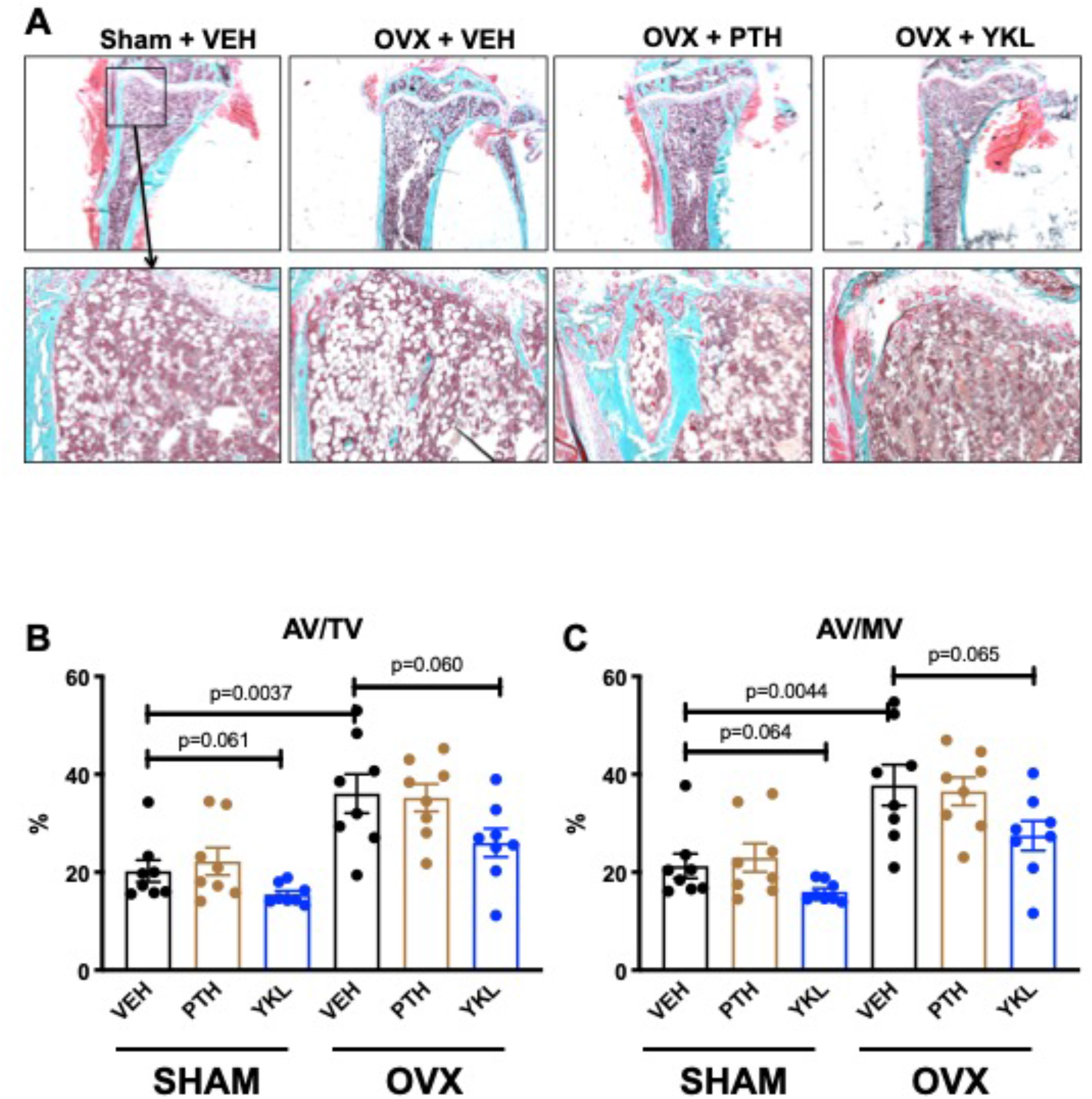
Effects of YKL-05-099 on marrow adipocytes in the proximal tibia. (A) Trichrome stained tibia sections for histomorphometry are shown. In 24 week old mice, many cells with adipocyte morphology are seen in the proximal tibial metaphysis. Qualitative reductions in adipocytes at this skeletal site are seen in YKL-05-099-treated mice. (B) Marrow adipocytes were quantified in a blinded manner. P values between groups were calculated by one way ANOVA followed by Dunnett’s correction. All p values less than 0.1 are shown. OVX surgery increases tissue space occupied by marrow adipocytes, and YKL-05-099 treatment tends to reduce this parameter. A high resolution version of this figure is available here: https://www.dropbox.com/s/g2t28largh4euvs/Figure%20S2.pdf?dl=0

**Supplemental Figure 3.**
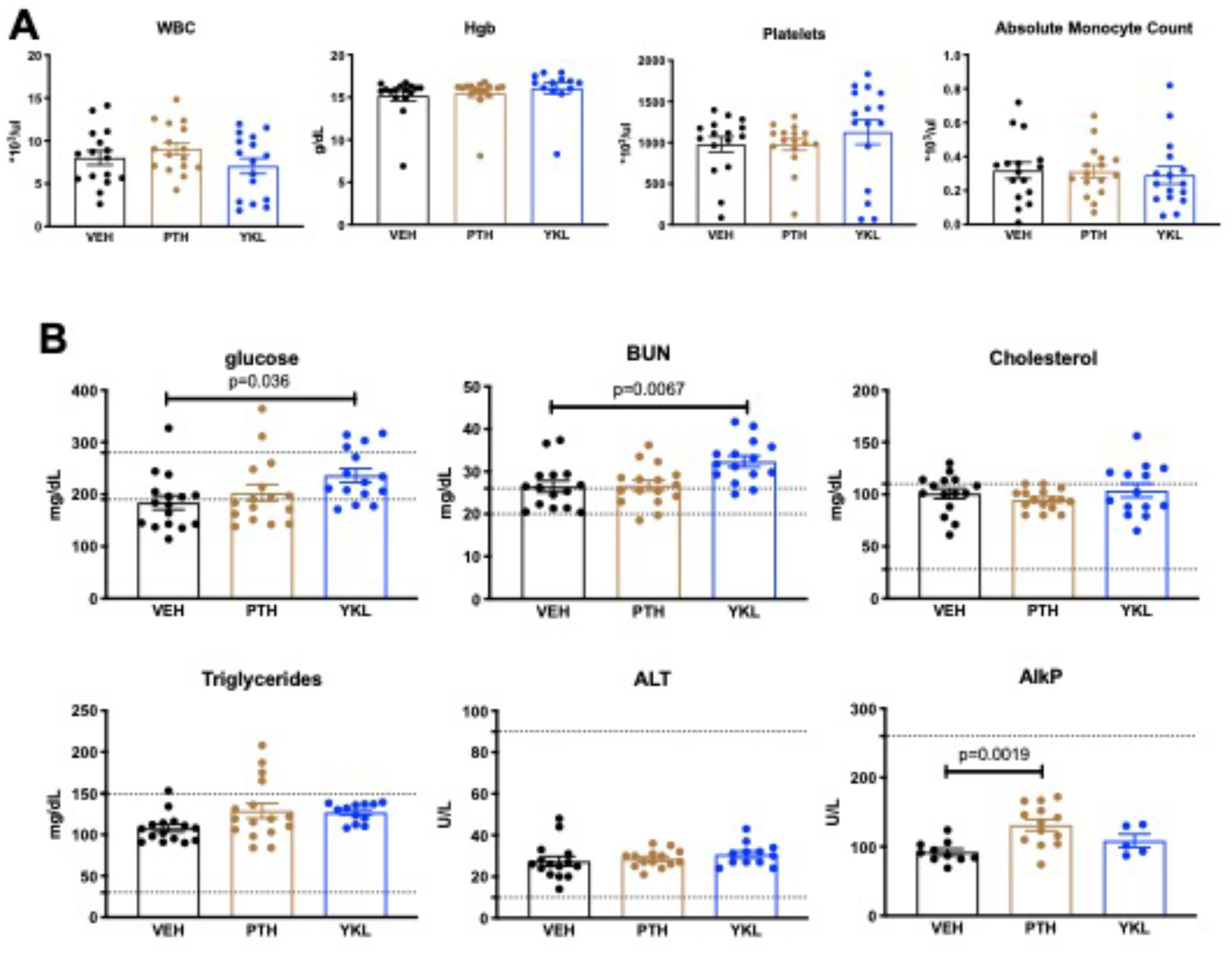
Effects of YKL-05-099 treatment on basic hematologic and serum parameters. 3 hour fasting blood was collected prior to sacrifice and analyzed for complete blood counts (A, WBC = white blood cells, Hgb = hemoglobin) and the indicated serum parameters (B, BUN = blood urea nitrogen, ALT = alanine transaminase, AlkP = alkaline phosphatase). For (B), the normal mouse reference ranges are shown in dotted horizontal lines. P values between groups were calculated by one way ANOVA followed by Dunnett’s correction. All p values less than 0.05 are shown. For these analyses, no effects of surgical intervention (sham versus ovariectomy) were noted. Therefore, mice are grouped based on drug treatment alone. No significant changes in hematologic parameters were noted. YKL-05-099 treatment led to statistically-significant increases in serum glucose and BUN. A high resolution version of this figure is available here: https://www.dropbox.com/s/306kbvt8xtg2z8h/Figure%20S3.pdf?dl=0

**Supplemental Figure 4.**
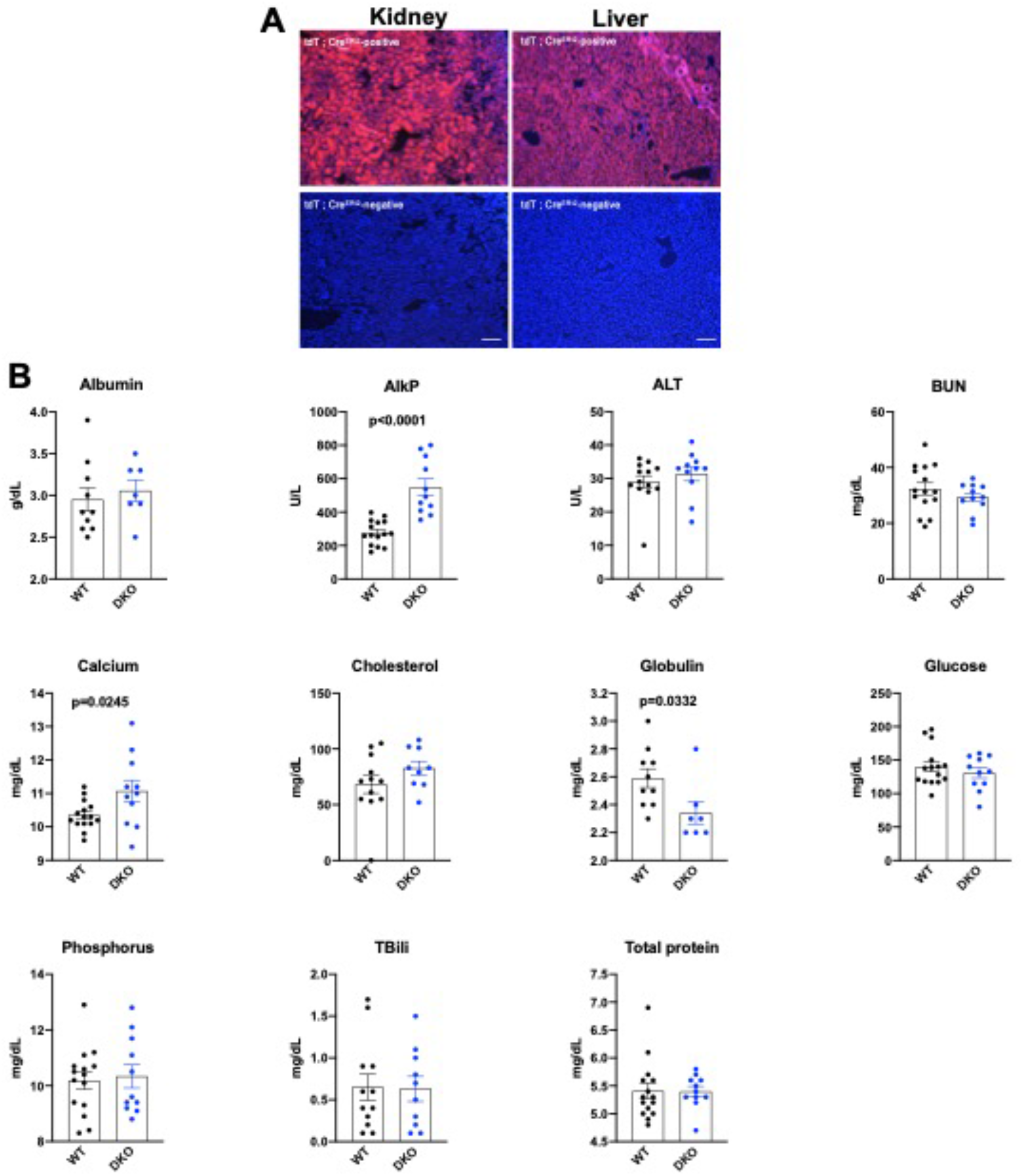
Efficacy and safety of global inducible Sik2/3 deletion. (A) Ubiquitin-Cre^ERt2^ mice were crossed to tdTomato reporter (Ai14) animals. 6 week old mice were treated with tamoxifen (1 mg by intraperitoneal injection, every other day, 3 doses total) and then sacrificed 7 days after the first tamoxifen injection. Cryosections of the kidney and liver were analyzed for tdTomato fluorescence. Bottom panels show kidney and liver sections from ubiquitin-Cre^ERt2^ mice that are negative for the Ai14 reporter allele. (B) *Sik2*^f/f^ ; *Sik3*^f/f^ (WT) and *Sik2*^f/f^ ; *Sik3*^f/f^ ; ubiquitin-Cre^ERt2^ (DKO) mice were treated with tamoxifen (1 mg by intraperitoneal injection, every other day, 3 doses total) starting at 6 weeks of age. 2 weeks later (8 weeks of age), fasting serum was collected for analysis of the indicated parameters. P values between groups were calculated by student’s t-test. All p values less than 0.05 are shown on individual graphs. *Sik2/3* deletion in this model led to statistically significant increases in AlkP and calcium, and slight reduction in globulin levels. Notably, BUN and glucose were not increased by tamoxifen-induced ubiquitous Sik2/3 deletion. A high resolution version of this figure is available here: https://www.dropbox.com/s/rbr7lk42e96avhz/Figure%20S4.pdf?dl=0

**Supplemental Figure 5.**
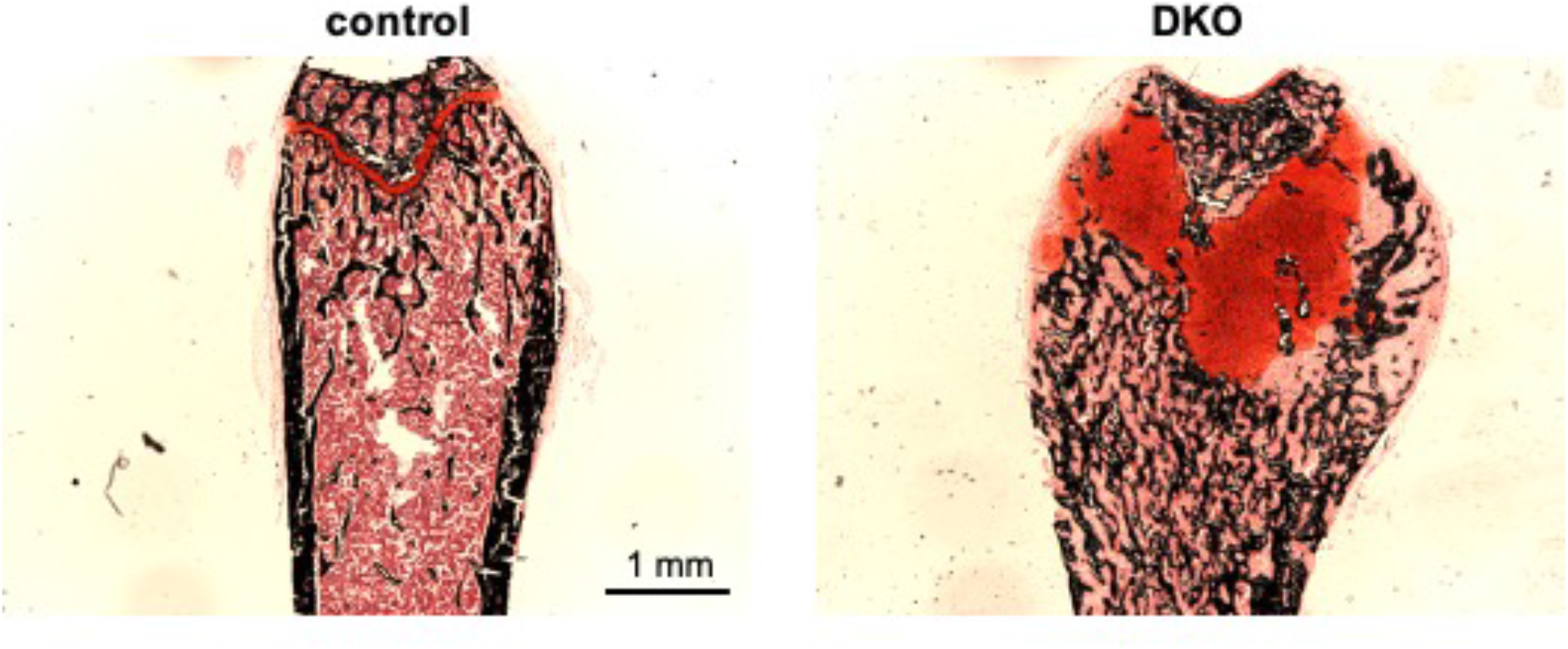
Growth plate defect induced by post-natal SIK2/3 deletion. 6 week old Sik2^f/f^ ; Sik3^f/f^ (WT) or Sik2^f/f^ ; Sik3^f/f^ ; ubiquitin-Cre^ERt2^ (DKO) mice were treated with tamoxifen (1 mg by intraperitoneal injection, every other day, 3 doses total) and then sacrificed 21 days after the first tamoxifen injection. Non-decalcified sections were obtained from the tibia which were stained with von Kossa (black) and safranin O (red). Dramatic growth plate expansion and disorganization is noted with inducible *Sik2/3* deletion. See also Figure 3E. A high resolution version of this figure is available here: https://www.dropbox.com/s/e8ws6mpcycbgqx8/Figure%20S5.pdf?dl=0

**Supplemental Figure 6.**
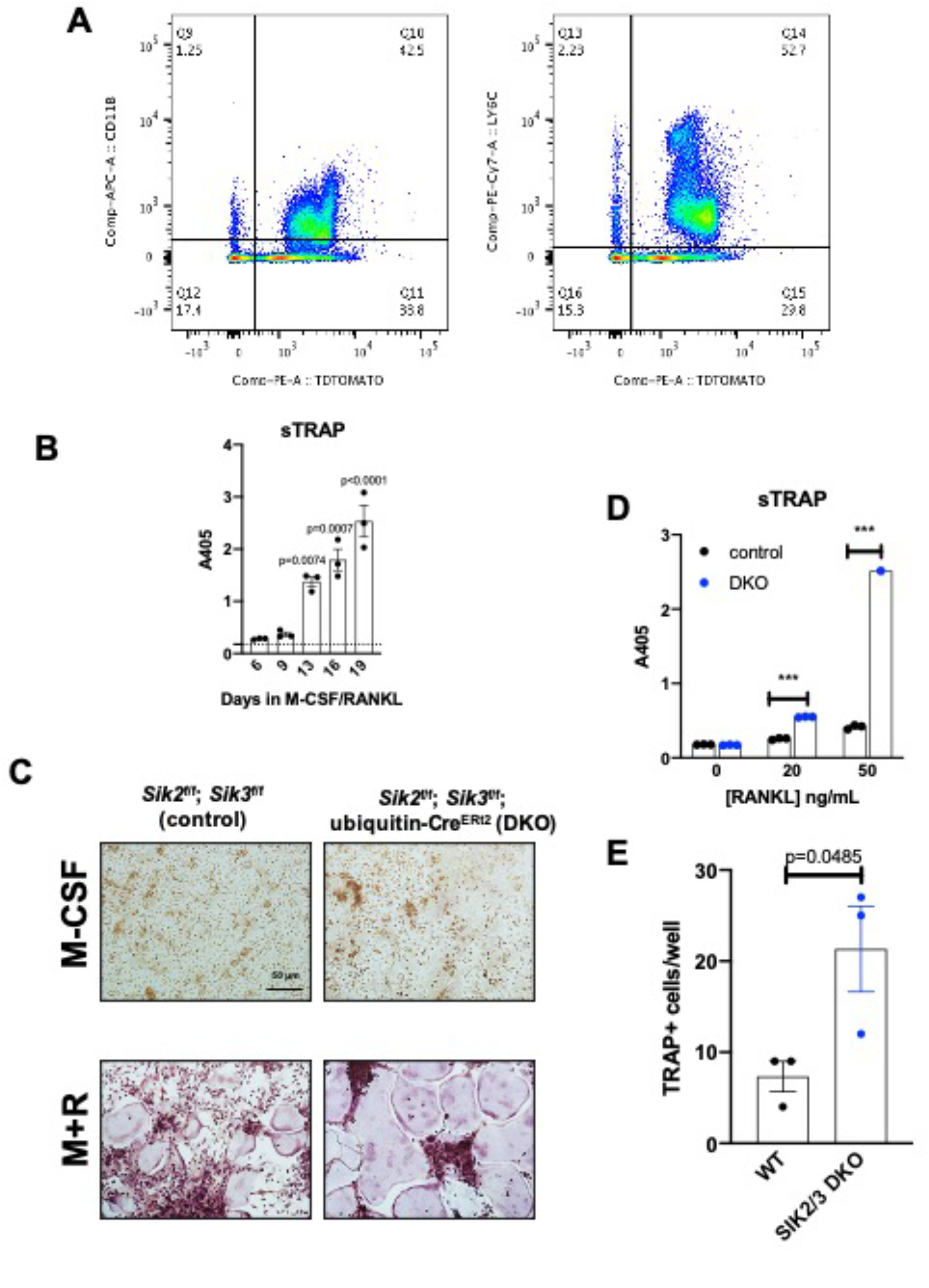
In vivo SIK2/3 deletion increases ex vivo osteoclast differentiation. (A) Ubiquitin-Cre^ERt2^ ; Ai14 reporter mice (tdTomato^LSL^) were treated with tamoxifen (1 mg by intraperitoneal injection, every other day, 3 doses total) starting at 6 weeks of age. Mice were sacrificed 2 weeks after the first tamoxifen dose and bone marrow cells were analyzed by flow cytometry. The majority (>90%) of myeloid lineage cells, as marked by CD11B (left) or LY6C (right), show evidence of Cre^ERt2^ activity as assessed by tdTomato protein. 97% of CD11B^+^ cells are tdTomato^+^, and 96% of LY6C^+^ cells are tdTomato^+^. (B) Bone marrow macrophages were grown in the presence of M-CSF plus RANKL for the indicated times. Culture supernatants were collected and secreted TRAP activity was measured over time. Robust osteoclast differentiation is noted after 10-14 days in these culture conditions. (C) Bone marrow cells were isolated from mice 2 weeks after in vivo tamoxifen administration as in (A). Bone marrow macrophages were isolated and grown in the presence of M-CSF alone (top) or M-CSF plus RANKL (M+R, bottom) followed by TRAP staining (purple). Cells isolated from DKO mice treated with tamoxifen in vivo show increased osteoclast differentiation. Scale bar = 50 µm (D) Culture supernatants from (C) were assessed for secreted TRAP activity. (E) TRAP-positive multinucleated cells from (C) treated with RANKL 50 ng/mL were quantified. A high resolution version of this figure is available here: https://www.dropbox.com/s/lskwrtk4nuxsyon/Figure%20S6.pdf?dl=0

**Supplemental Figure 7.**
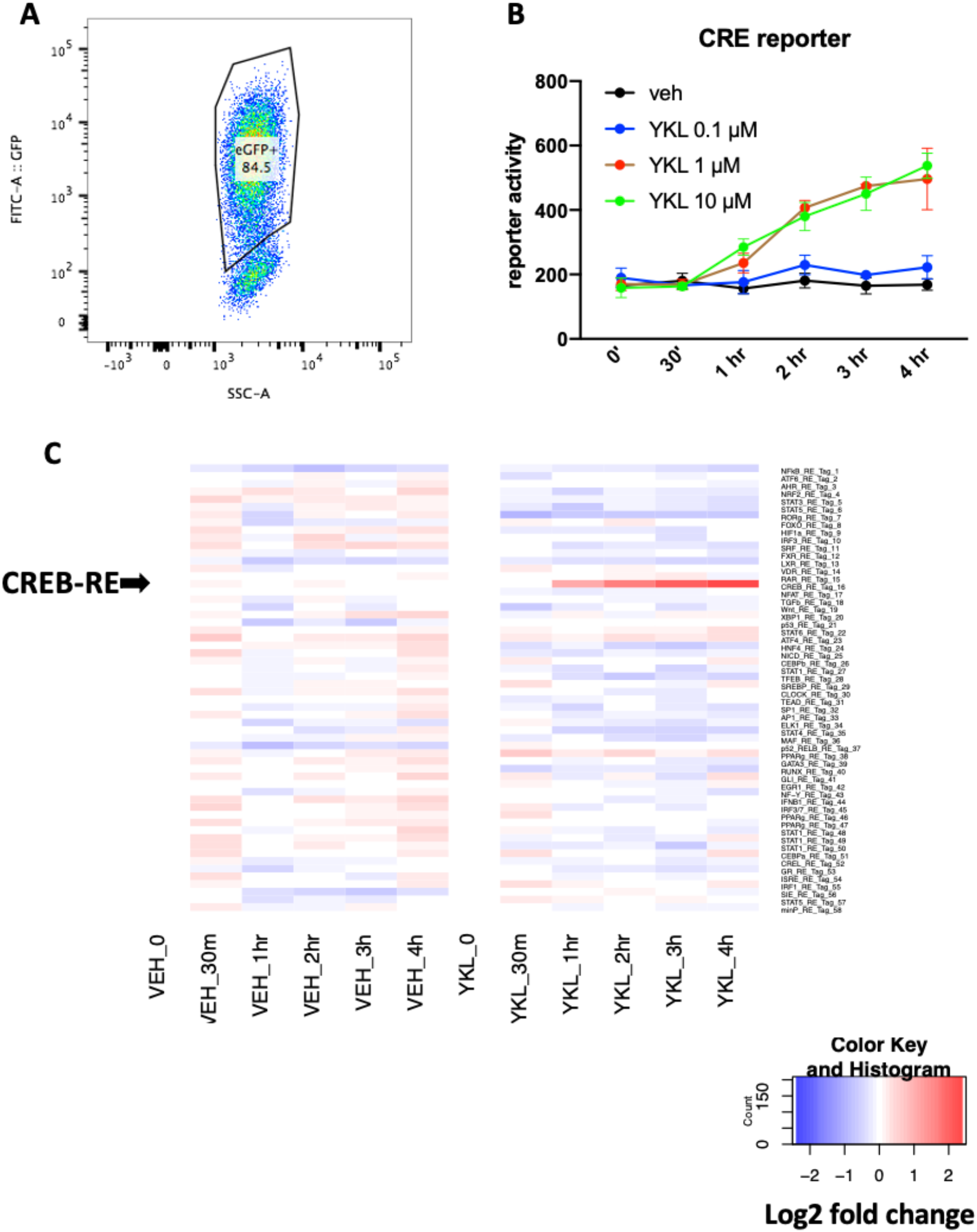
Parallel reporter assay (TF-seq) was used to determine the effects of YKL-05-099 on 58 reporter synthetic reporter elements. (A) Ocy454 cells were infected with TF-seq lentiviral particles which co-express GFP using spin infection in the presence of polybrene (2 µg/ml). Infected cells were expanded and then subjected to flow cytometry. GFP+ cells were sorted for subsequent experiments. (B) Sorted GFP+ TF-seq Ocy454 cells were treated with the indicated dose of YKL-05-099 or DMSO (vehicle) control for the indicated times. Cells were lysed and reporter element (RE) for each of the 58 elements was measured as detailed in the methods. Of the 58 elements tested, only CRE activity was significantly regulated by YKL-05-099 treatment. Panel B shows the effects of YKL-05-099 on CRE activity. (C) Heat map showing activity of all 58 reporter elements in response to vehicle or 10 µM YKL-05-099. Each row represents a distinct reporter element. Arrowhead (left) shows the row that corresponds to the CRE element. Color code shows log2 fold change relative to time zero. A high resolution version of this figure is available here: https://www.dropbox.com/s/g6i4nwtwphrlnym/Figure%20S7.pdf?dl=0

**Supplemental Figure 8.**
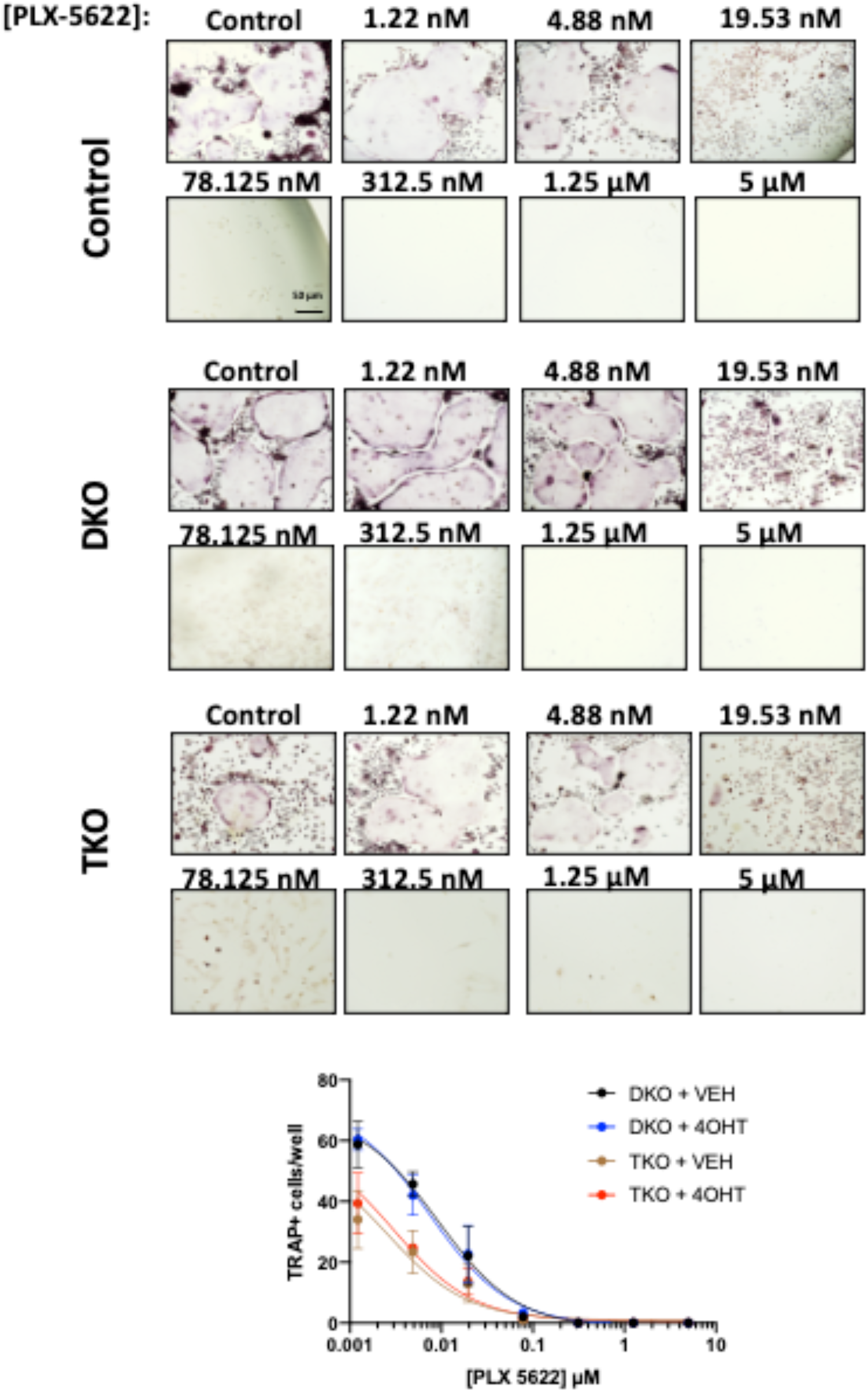
PLX-5622 blocks osteoclast differentiation in SIK mutant cells. Bone marrow macrophages from ubiquitin-Cre^ERt2^ ; SIK2/3 (DKO) or ubiquitin-Cre^ERt2^ ; SIK1/2/3 (TKO) floxed mice were treated plus/minus 4-OHT and then were grown in the presence of M-CSF/RANKL plus the indicated doses of PLX-5622 for 12 days followed by TRAP staining. Representative photomicrographs from the indicated cells and drug concentration are shown. Scale bar = 50 µm. Bottom, quantification of TRAP+ multinucleated cells. For each condition, n=3 wells (96 well plate) were analyzed, error bars represent mean ± SD. PLX-5622 reduced numbers of TRAP+ multinucleated cells irrespective of SIK genotype. A high resolution version of this figure is available here: https://www.dropbox.com/s/grvou68j9to1uwo/Figure%20S8.pdf?dl=0

**Supplemental Figure 9.**
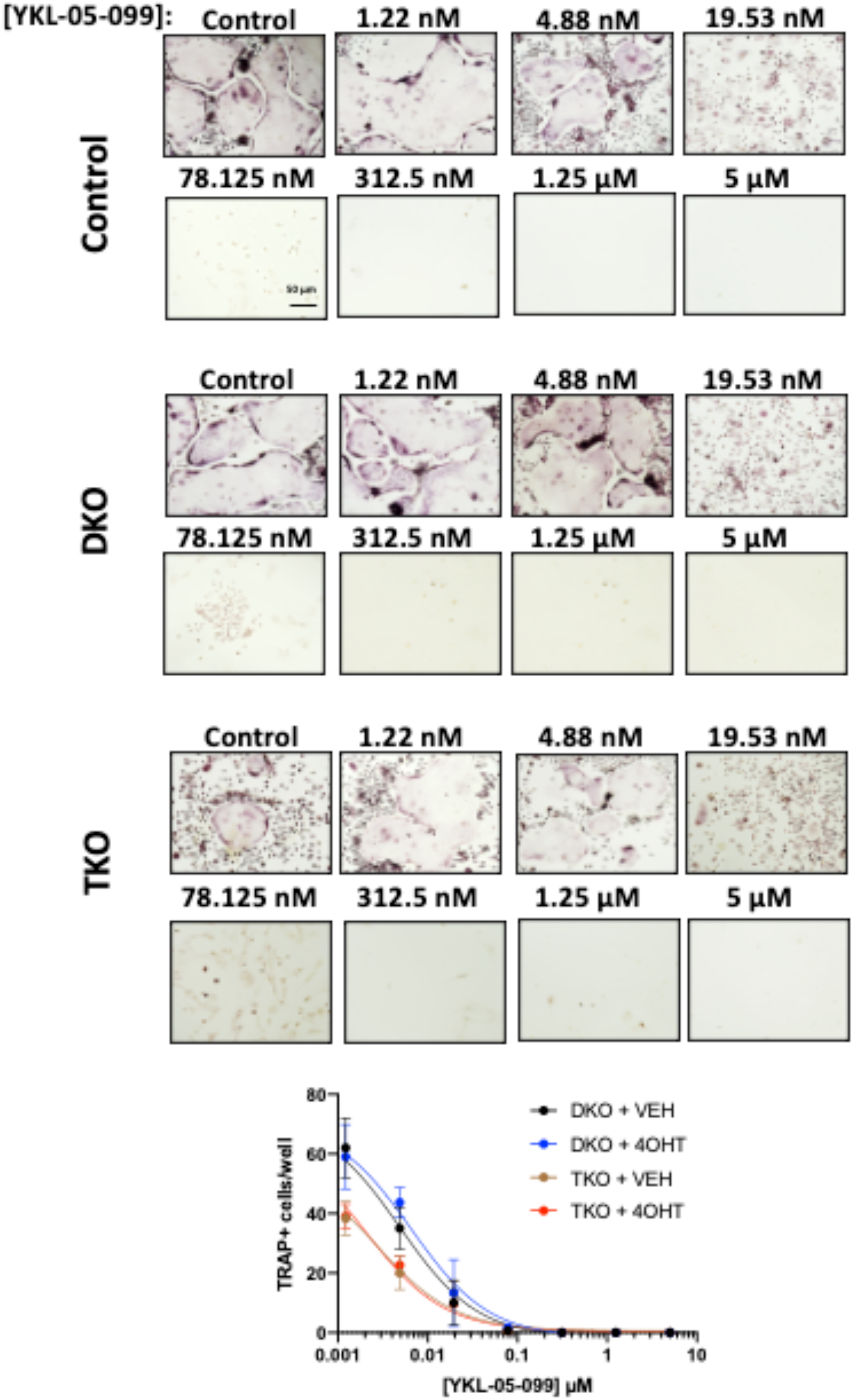
YKL-05-099 blocks osteoclast differentiation in SIK mutant cells. Bone marrow macrophages from ubiquitin-Cre^ERt2^ ; SIK2/3 (DKO) or ubiquitin-Cre^ERt2^ ; SIK1/2/3 (TKO) floxed mice were treated plus/minus 4-OHT and then were grown in the presence of M-CSF/RANKL plus the indicated doses of YKL-05-099 for 12 days followed by TRAP staining. Representative photomicrographs from the indicated cells and drug concentration are shown. Scale bar = 50 µm. Bottom, quantification of TRAP+ multinucleated cells. For each condition, n=3 wells (96 well plate) were analyzed, error bars represent mean ± SD. YKL-05-099 reduced numbers of TRAP+ multinucleated cells irrespective of SIK genotype. A high resolution version of this figure is available here: https://www.dropbox.com/s/qfivnhwf7o59dxf/Figure%20S9.pdf?dl=0

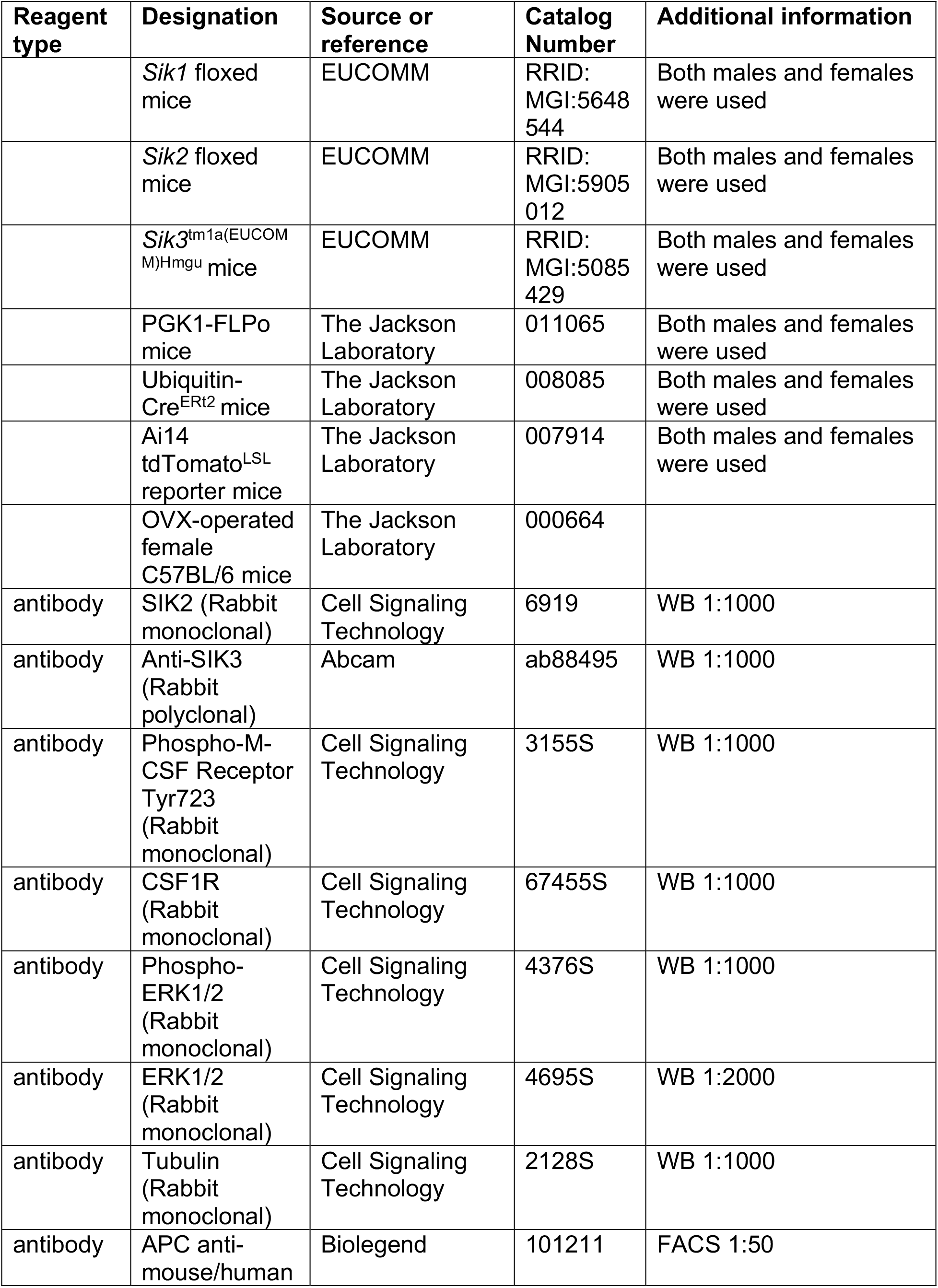
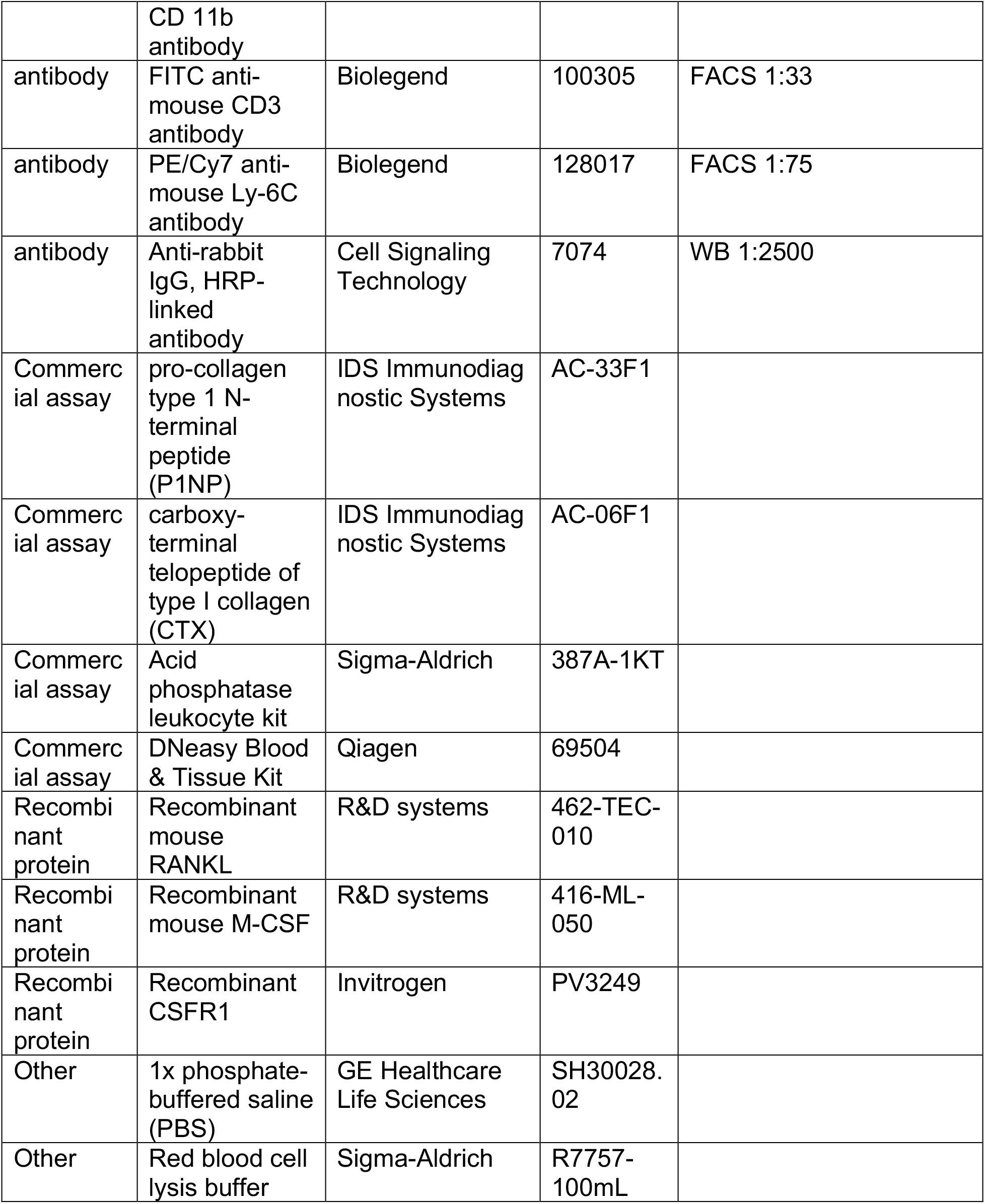
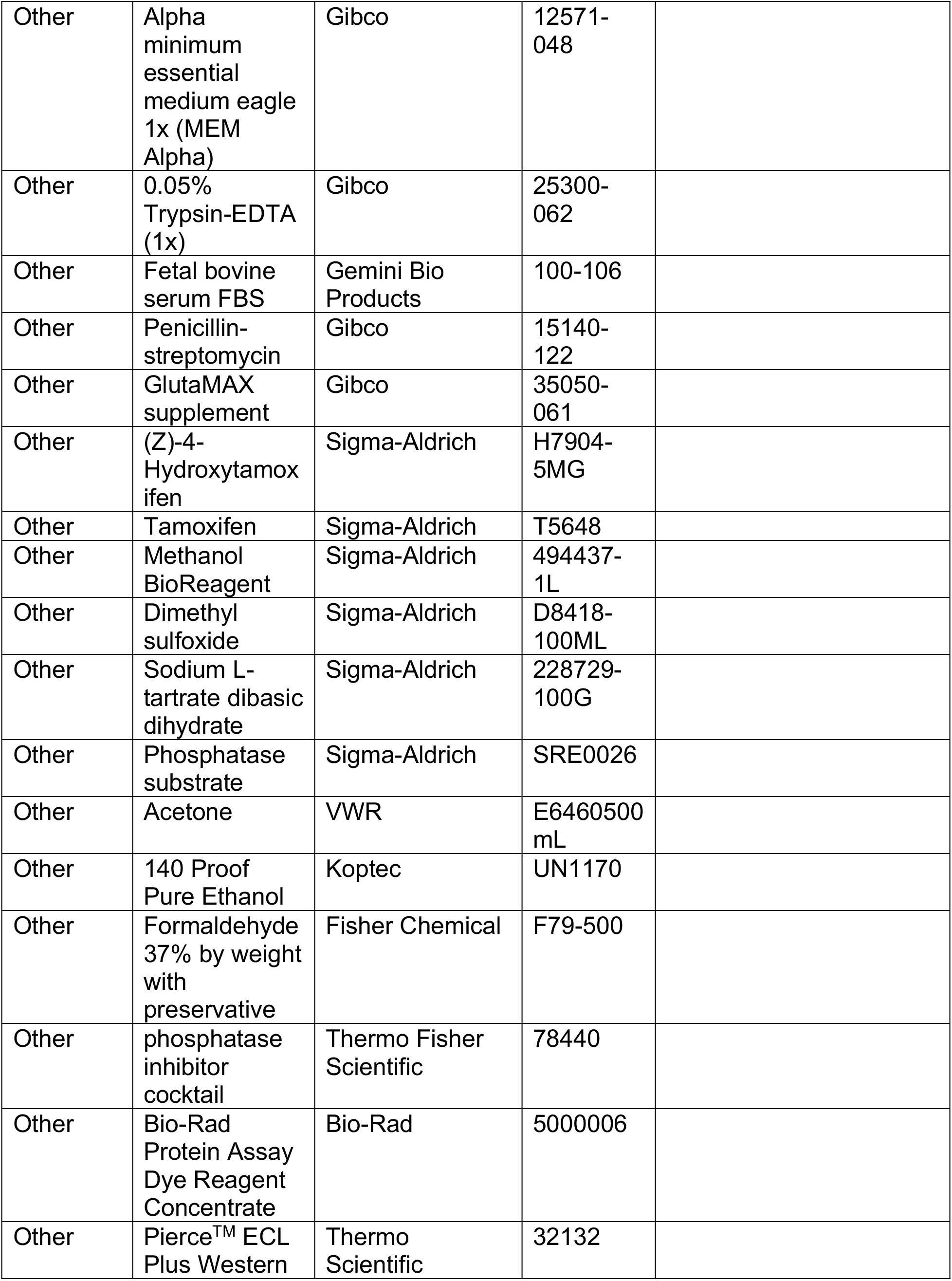
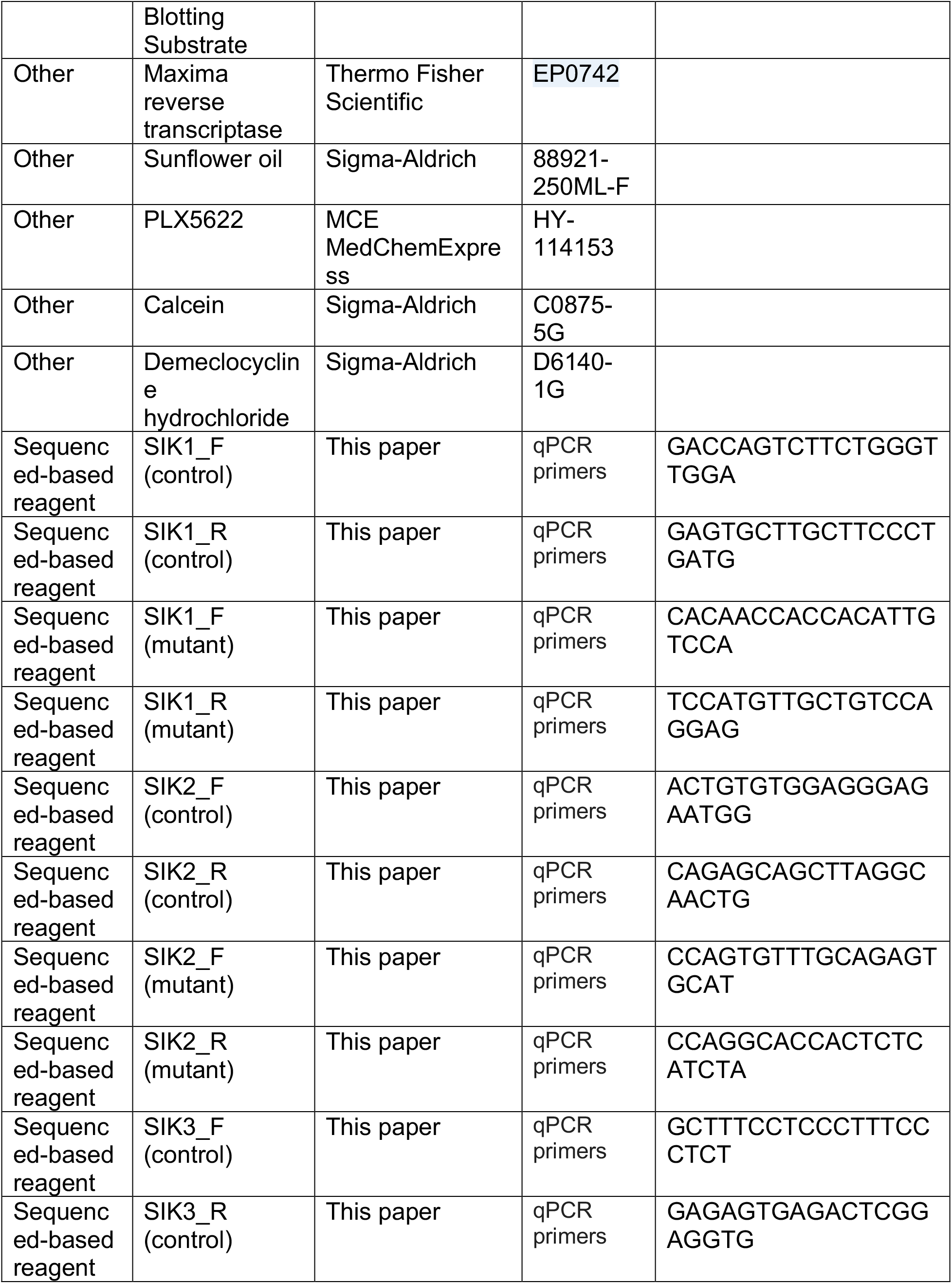
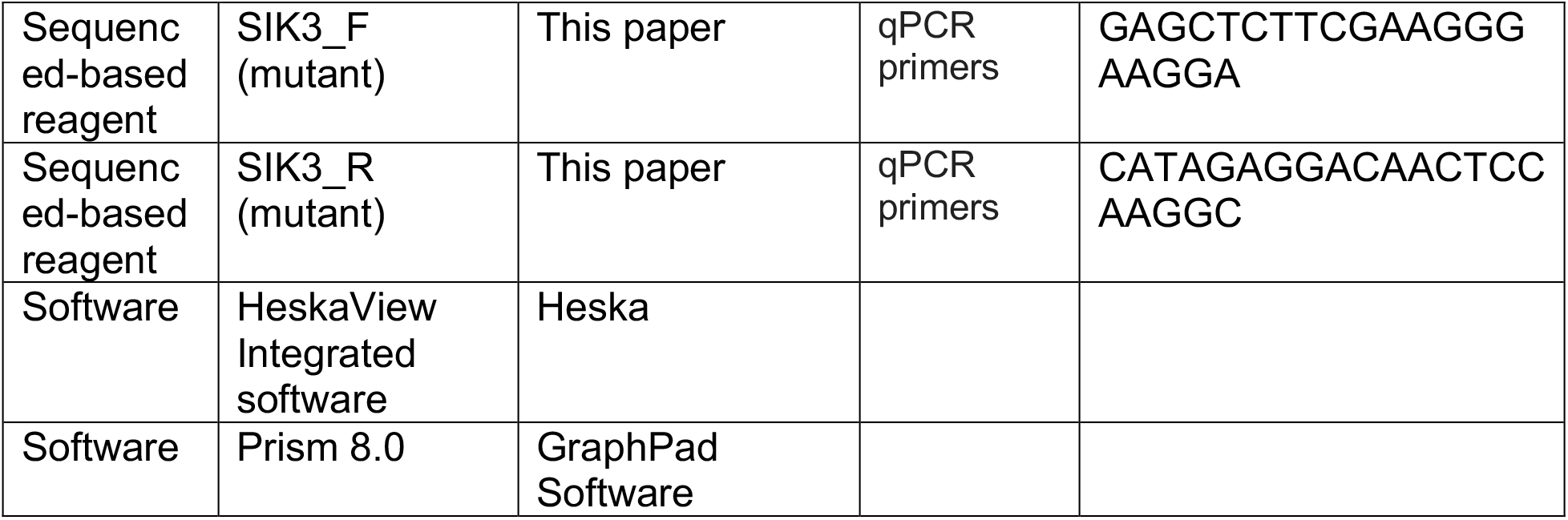

## References

1. Harvey N, Dennison E, and Cooper C. Osteoporosis: impact on health and economics. Nature reviews Rheumatology. 2010;6(2):99–105.

2. Zaidi M. Skeletal remodeling in health and disease. Nature medicine. 2007;13(7):791–801.

3. Dallas SL, Prideaux M, and Bonewald LF. The osteocyte: an endocrine cell … and more. Endocrine reviews. 2013;34(5):658–90.

4. Compston JE, McClung MR, and Leslie WD. Osteoporosis. Lancet. 2019;393(10169):364–76.

5. Estell EG, and Rosen CJ. Emerging insights into the comparative effectiveness of anabolic therapies for osteoporosis. Nature reviews Endocrinology. 2021;17(1):31–46.

6. Bilezikian JP. Combination anabolic and antiresorptive therapy for osteoporosis: opening the anabolic window. Current osteoporosis reports. 2008;6(1):24–30.

7. Wein MN. Parathyroid Hormone Signaling in Osteocytes. JBMR Plus. 2018;2(1):22–30.

8. Sakamoto K, Bultot L, and Goransson O. The Salt-Inducible Kinases: Emerging Metabolic Regulators. Trends Endocrinol Metab. 2018;29(12):827–40.

9. Wein MN, Foretz M, Fisher DE, Xavier RJ, and Kronenberg HM. Salt-Inducible Kinases: Physiology, Regulation by cAMP, and Therapeutic Potential. Trends Endocrinol Metab. 2018.

10. Wein MN, Liang Y, Goransson O, Sundberg TB, Wang J, Williams EA, et al. SIKs control osteocyte responses to parathyroid hormone. Nature communications. 2016;7:13176.

11. Nishimori S, O’Meara MJ, Castro CD, Noda H, Cetinbas M, da Silva Martins J, et al. Salt-inducible kinases dictate parathyroid hormone 1 receptor action in bone development and remodeling. J Clin Invest. 2019.

12. Sundberg TB, Liang Y, Wu H, Choi HG, Kim ND, Sim T, et al. Development of Chemical Probes for Investigation of Salt-Inducible Kinase Function in Vivo. ACS chemical biology. 2016.

13. Sims NA, and Martin TJ. Osteoclasts Provide Coupling Signals to Osteoblast Lineage Cells Through Multiple Mechanisms. Annu Rev Physiol. 2020;82:507–29.

14. Tarumoto Y, Lin S, Wang J, Milazzo JP, Xu Y, Lu B, et al. Salt-Inducible Kinase inhibition suppresses acute myeloid leukemia progression in vivo. Blood. 2019.

15. Dar AC, Das TK, Shokat KM, and Cagan RL. Chemical genetic discovery of targets and anti-targets for cancer polypharmacology. Nature. 2012;486(7401):80–4.

16. Ferguson FM, and Gray NS. Kinase inhibitors: the road ahead. Nature reviews Drug discovery. 2018;17(5):353–77.

17. Mun SH, Park PSU, and Park-Min KH. The M-CSF receptor in osteoclasts and beyond. Exp Mol Med. 2020;52(8):1239–54.

18. Roschger P, Paschalis EP, Fratzl P, and Klaushofer K. Bone mineralization density distribution in health and disease. Bone. 2008;42(3):456–66.

19. de Paula FJA, and Rosen CJ. Marrow Adipocytes: Origin, Structure, and Function. Annu Rev Physiol. 2020;82:461–84.

20. Fan Y, Hanai JI, Le PT, Bi R, Maridas D, DeMambro V, et al. Parathyroid Hormone Directs Bone Marrow Mesenchymal Cell Fate. Cell metabolism. 2017;25(3):661–72.

21. Balani DH, Ono N, and Kronenberg HM. Parathyroid hormone regulates fates of murine osteoblast precursors in vivo. J Clin Invest. 2017;127(9):3327–38.

22. Yang M, Arai A, Udagawa N, Zhao L, Nishida D, Murakami K, et al. Parathyroid Hormone Shifts Cell Fate of a Leptin Receptor-Marked Stromal Population from Adipogenic to Osteoblastic Lineage. Journal of bone and mineral research : the official journal of the American Society for Bone and Mineral Research. 2019;34(10):1952–63.

23. Maridas DE, Rendina-Ruedy E, Helderman RC, DeMambro VE, Brooks D, Guntur AR, et al. Progenitor recruitment and adipogenic lipolysis contribute to the anabolic actions of parathyroid hormone on the skeleton. FASEB J. 2019;33(2):2885–98.

24. Tratwal J, Labella R, Bravenboer N, Kerckhofs G, Douni E, Scheller EL, et al. Reporting Guidelines, Review of Methodological Standards, and Challenges Toward Harmonization in Bone Marrow Adiposity Research. Report of the Methodologies Working Group of the International Bone Marrow Adiposity Society. Frontiers in endocrinology. 2020;11:65.

25. Patel K, Foretz M, Marion A, Campbell DG, Gourlay R, Boudaba N, et al. The LKB1-salt-inducible kinase pathway functions as a key gluconeogenic suppressor in the liver. Nature communications. 2014;5:4535.

26. Ruzankina Y, Pinzon-Guzman C, Asare A, Ong T, Pontano L, Cotsarelis G, et al. Deletion of the developmentally essential gene ATR in adult mice leads to age-related phenotypes and stem cell loss. Cell stem cell. 2007;1(1):113–26.

27. Sasagawa S, Takemori H, Uebi T, Ikegami D, Hiramatsu K, Ikegawa S, et al. SIK3 is essential for chondrocyte hypertrophy during skeletal development in mice. Development. 2012;139(6):1153–63.

28. MD ZXMCMJSPMSMSLLPADPAJvWPJRMBS. Low Dose Tamoxifen Induces Significant Bone Formation in Mice. JBMR PLUS. 2020.

29. Lombardi MS, Gillieron C, Berkelaar M, and Gabay C. Salt-inducible kinases (SIK) inhibition reduces RANKL-induced osteoclastogenesis. PloS one. 2017;12(10):e0185426.

30. Chen K, Ng PY, Chen R, Hu D, Berry S, Baron R, et al. Sfrp4 repression of the Ror2/Jnk cascade in osteoclasts protects cortical bone from excessive endosteal resorption. Proceedings of the National Academy of Sciences of the United States of America. 2019;116(28):14138–43.

31. Nixon M, Stewart-Fitzgibbon R, Fu J, Akhmedov D, Rajendran K, Mendoza-Rodriguez MG, et al. Skeletal muscle salt inducible kinase 1 promotes insulin resistance in obesity. Molecular metabolism. 2016;5(1):34–46.

32. Ricarte FR, Le Henaff C, Kolupaeva VG, Gardella TJ, and Partridge NC. Parathyroid hormone (1-34) and its analogs differentially modulate osteoblastic RANKL expression via PKA/PP1/PP2A and SIK2/SIK3-CRTC3 signaling. J BiolD ChemD. 2018.

33. O’Connell DJ, Kolde R, Sooknah M, Graham DB, Sundberg TB, Latorre I, et al. Simultaneous Pathway Activity Inference and Gene Expression Analysis Using RNA Sequencing. Cell systems. 2016;2(5):323–34.

34. Spatz JM, Wein MN, Gooi JH, Qu Y, Garr JL, Liu S, et al. The Wnt Inhibitor Sclerostin Is Up-regulated by Mechanical Unloading in Osteocytes in Vitro. J BiolD ChemD. 2015;290(27):16744–58.

35. Wein MN, Spatz J, Nishimori S, Doench J, Root D, Babij P, et al. HDAC5 controls MEF2C-driven sclerostin expression in osteocytes. Journal of bone and mineral research : the official journal of the American Society for Bone and Mineral Research. 2015;30(3):400–11.

36. Altarejos JY, and Montminy M. CREB and the CRTC co-activators: sensors for hormonal and metabolic signals. Nature reviews Molecular cell biology. 2011;12(3):141–51.

37. Ross FP, and Teitelbaum SL. alphavbeta3 and macrophage colony-stimulating factor: partners in osteoclast biology. Immunol Rev. 2005;208:88–105.

38. Manning G, Whyte DB, Martinez R, Hunter T, and Sudarsanam S. The protein kinase complement of the human genome. Science. 2002;298(5600):1912–34.

39. Ung PM, and Schlessinger A. DFGmodel: predicting protein kinase structures in inactive states for structure-based discovery of type-II inhibitors. ACS chemical biology. 2015;10(1):269–78.

40. Roskoski R, Jr. Classification of small molecule protein kinase inhibitors based upon the structures of their drug-enzyme complexes. Pharmacol Res. 2016;103:26–48.

41. Ung PM, Rahman R, and Schlessinger A. Redefining the Protein Kinase Conformational Space with Machine Learning. Cell Chem Biol. 2018;25(7):916–24 e2.

42. Rahman R, Ung PM, and Schlessinger A. KinaMetrix: a web resource to investigate kinase conformations and inhibitor space. Nucleic Acids Res. 2019;47(D1):D361–D6.

43. Dar AC, and Shokat KM. The evolution of protein kinase inhibitors from antagonists to agonists of cellular signaling. Annu Rev Biochem. 2011;80:769–95.

44. Bourette RP, Myles GM, Choi JL, and Rohrschneider LR. Sequential activation of phoshatidylinositol 3-kinase and phospholipase C-gamma2 by the M-CSF receptor is necessary for differentiation signaling. The EMBO journal. 1997;16(19):5880–93.

45. Tran DD, Saran S, Dittrich-Breiholz O, Williamson AJ, Klebba-Farber S, Koch A, et al. Transcriptional regulation of immediate-early gene response by THOC5, a member of mRNA export complex, contributes to the M-CSF-induced macrophage differentiation. Cell Death Dis. 2013;4:e879.

46. Spangenberg E, Severson PL, Hohsfield LA, Crapser J, Zhang J, Burton EA, et al. Sustained microglial depletion with CSF1R inhibitor impairs parenchymal plaque development in an Alzheimer’s disease model. Nature communications. 2019;10(1):3758.

47. McClung MR, Grauer A, Boonen S, Bolognese MA, Brown JP, Diez-Perez A, et al. Romosozumab in postmenopausal women with low bone mineral density. N Engl J Med. 2014;370(5):412–20.

48. Fixen C, and Tunoa J. Romosozumab: a Review of Efficacy, Safety, and Cardiovascular Risk. Current osteoporosis reports. 2021.

49. Zhou J, Alfraidi A, Zhang S, Santiago-O’Farrill JM, Yerramreddy Reddy VK, Alsaadi A, et al. A Novel Compound ARN-3236 Inhibits Salt-Inducible Kinase 2 and Sensitizes Ovarian Cancer Cell Lines and Xenografts to Paclitaxel. Clinical cancer research : an official journal of the American Association for Cancer Research. 2017;23(8):1945–54.

50. Darling NJ, Toth R, Arthur JS, and Clark K. Inhibition of SIK2 and SIK3 during differentiation enhances the anti-inflammatory phenotype of macrophages. The Biochemical journal. 2017;474(4):521–37.

51. Berggreen C, Henriksson E, Jones HA, Morrice N, and Goransson O. cAMP-elevation mediated by beta-adrenergic stimulation inhibits salt-inducible kinase (SIK) 3 activity in adipocytes. Cell Signal. 2012;24(9):1863–71.

52. Henriksson E, Jones HA, Patel K, Peggie M, Morrice N, Sakamoto K, et al. The AMPK-related kinase SIK2 is regulated by cAMP via phosphorylation at Ser358 in adipocytes. The Biochemical journal. 2012;444(3):503–14.

53. Scheller EL, Khandaker S, Learman BS, Cawthorn WP, Anderson LM, Pham HA, et al. Bone marrow adipocytes resist lipolysis and remodeling in response to beta-adrenergic stimulation. Bone. 2019;118:32–41.

54. Murshed M, and McKee MD. Molecular determinants of extracellular matrix mineralization in bone and blood vessels. Curr Opin Nephrol Hypertens. 2010;19(4):359–65.

55. Zhang J, Yang PL, and Gray NS. Targeting cancer with small molecule kinase inhibitors. Nat Rev Cancer. 2009;9(1):28–39.

56. Liu Y, Shah K, Yang F, Witucki L, and Shokat KM. A molecular gate which controls unnatural ATP analogue recognition by the tyrosine kinase v-Src. Bioorg Med Chem. 1998;6(8):1219–26.

57. Azam M, Seeliger MA, Gray NS, Kuriyan J, and Daley GQ. Activation of tyrosine kinases by mutation of the gatekeeper threonine. Nat Struct Mol Biol. 2008;15(10):1109–18.

58. Klaeger S, Heinzlmeir S, Wilhelm M, Polzer H, Vick B, Koenig PA, et al. The target landscape of clinical kinase drugs. Science. 2017;358(6367).

59. Hanson SM, Georghiou G, Thakur MK, Miller WT, Rest JS, Chodera JD, et al. What Makes a Kinase Promiscuous for Inhibitors? Cell Chem Biol. 2019;26(3):390–9 e5.

60. Grassi F, Tyagi AM, Calvert JW, Gambari L, Walker LD, Yu M, et al. Hydrogen Sulfide Is a Novel Regulator of Bone Formation Implicated in the Bone Loss Induced by Estrogen Deficiency. Journal of bone and mineral research : the official journal of the American Society for Bone and Mineral Research. 2016;31(5):949–63.

61. Bouxsein ML, Boyd SK, Christiansen BA, Guldberg RE, Jepsen KJ, and Muller R. Guidelines for assessment of bone microstructure in rodents using micro-computed tomography. Journal of bone and mineral research : the official journal of the American Society for Bone and Mineral Research. 2010;25(7):1468–86.

62. Dempster DW, Compston JE, Drezner MK, Glorieux FH, Kanis JA, Malluche H, et al. Standardized nomenclature, symbols, and units for bone histomorphometry: a 2012 update of the report of the ASBMR Histomorphometry Nomenclature Committee. Journal of bone and mineral research : the official journal of the American Society for Bone and Mineral Research. 2013;28(1):2–17.

63. Schneider CA, Rasband WS, and Eliceiri KW. NIH Image to ImageJ: 25 years of image analysis. Nature methods. 2012;9(7):671–5.

64. van ’t Hof RJ, Rose L, Bassonga E, and Daroszewska A. Open source software for semi-automated histomorphometry of bone resorption and formation parameters. Bone. 2017;99:69–79.

65. Roschger P, Fratzl P, Eschberger J, and Klaushofer K. Validation of quantitative backscattered electron imaging for the measurement of mineral density distribution in human bone biopsies. Bone. 1998;23(4):319–26.

66. Livak KJ, and Schmittgen TD. Analysis of relative gene expression data using real-time quantitative PCR and the 2(-Delta Delta C(T)) Method. Methods. 2001;25(4):402–8.

67. Modi V, and Dunbrack RL, Jr. A Structurally-Validated Multiple Sequence Alignment of 497 Human Protein Kinase Domains. Scientific reports. 2019;9(1):19790.

68. Camacho C, Coulouris G, Avagyan V, Ma N, Papadopoulos J, Bealer K, et al. BLAST+: architecture and applications. BMC Bioinformatics. 2009;10:421.

69. Sali A, and Blundell TL. Comparative protein modelling by satisfaction of spatial restraints. JD MolD BiolD. 1993;234(3):779–815.

70. Durrant JD, Votapka L, Sorensen J, and Amaro RE. POVME 2.0: An Enhanced Tool for Determining Pocket Shape and Volume Characteristics. Journal of chemical theory and computation. 2014;10(11):5047–56.

71. Friesner RA, Banks JL, Murphy RB, Halgren TA, Klicic JJ, Mainz DT, et al. Glide: a new approach for rapid, accurate docking and scoring. 1. Method and assessment of docking accuracy. J Med Chem. 2004;47(7):1739–49.

72. Harder E, Damm W, Maple J, Wu C, Reboul M, Xiang JY, et al. OPLS3: A Force Field Providing Broad Coverage of Drug-like Small Molecules and Proteins. Journal of chemical theory and computation. 2016;12(1):281–96.

